# Sidechain chemistry-encoded solid/liquid phase transitions of condensates

**DOI:** 10.1101/2024.09.16.613107

**Authors:** Feipeng Chen, Yongxu Han, Xiufeng Li, Wei Guo, Changjin Wu, Jiang Xia, Xiangze Zeng, Ho Cheung Shum

**Affiliations:** Department of Mechanical Engineering, The University of Hong Kong, Pokfulam Road, Hong Kong, China; Department of Chemistry, The Chinese University of Hong Kong, Shatin, Hong Kong, China; Department of Physics, Hong Kong Baptist University, Kowloon Tong, Hong Kong, China; Advanced Biomedical Instrumentation Centre, Hong Kong Science Park, Shatin, New Territories, Hong Kong (SAR) 999077, China; College of Food Science and Technology, Nanjing Agricultural University, Nanjing, 210095, Jiangsu, China

## Abstract

Nature effectively leverages multivalent interactions among fundamental building blocks in solvents to create remarkable materials for various purposes. One prominent example is the formation of biomolecular condensates through the phase separation of proteins and nucleic acids. In particular, these condensates play crucial roles in regulating cellular functions and constructing natural materials. During the phase separation, solvents not only provide liquid environments for solvating molecules but play crucial roles in affecting the material properties of condensates. However, it remains controversial in the literature that alcohol molecules, as one type of solvents, can solidify some condensates while also melting others, leading to liquid-to-solid phase transition (LSPT) or solid-to-liquid phase transitions (SLPT), respectively. The mechanism underlying the alcohol-induced solid/liquid phase transitions of condensates remains poorly understood. Here, we combine systematic experimental characterizations with molecular dynamics simulations to demonstrate that the phase transitions of condensates depend on their sidechain chemistry and dominant molecular interactions. Specifically, “hydrophilic” condensates, which consist of many charged sidechains, undergo LSPT by adding alcohols due to strengthened electrostatic interactions. In contrast, “hydrophobic” condensates comprised of abundant aromatic sidechains undergo SLPT with the addition of alcohols because of weakened cation-π and π-π interactions. Importantly, these findings are generally applicable for predicting phase transitions of a wide range of condensates formed by synthetic polyelectrolytes and intrinsically disordered proteins based on their sidechain hydrophobicity or amino acid compositions. Our work not only reconciles a conundrum in the literature but provides a fundamental framework for understanding the responsiveness of condensates to environmental stimuli. These insights are instrumental for developing therapeutic drugs to treat pathological aggregates and engineering stimuli-responsive biomaterials from the perspective of sidechain chemistry and molecular interactions.

## Introduction

Living materials form without human guidance, yet exhibit a wide range of properties that are often superior to most synthetic materials. Their formation usually relies on a bottom-up approach based on molecular interactions among building blocks within solvents [1]. For example, membrane-less biomolecular condensates are formed through multivalent interactions, such as electrostatic interactions, cation-π and π-π interactions, between macromolecules and small molecules within biological or synthetic systems [2, 3]. These condensates have attracted increasing attentions due to their crucial roles in regulating biological functions and in assembling remarkable materials observed in various living organisms [4, 5]. In these scenarios, the molecular interactions and physical properties of condensates can be dynamically tuned by varying solvent conditions, such as alcohols, salts, and temperature, which in turn affect their functions [6–8]. Notably, the phase transition of condensates from liquid droplets to more solid-like aggregates, known as the liquid-to-solid phase transition (LSPT), is suggested to trigger the pathogenesis of neurodegenerative diseases and cancers [9, 10]. LSPT could also provide structural functions for condensates, such as forming strong underwater adhesives by marine animals [11, 12]. However, how solvents affect the molecular interactions and thus the materials properties of condensates remains poorly understood; addressing this question could aid in the design of therapeutic drugs for treating pathological condensates and engineering synthetic materials for a plethora of applications [13–15].

This challenge is highlighted by an unresolved conundrum in the literature that shows inconsistent experimental results regarding the impact of alcohol molecules on the phase transition of condensates (summarized in Supplementary Table 1). For example, the introduction of ethanol was observed to enhance the aggregation of pathologically relevant proteins by crystallizing intermediate filament protein keratin-8 (KRT8)[16] and promoting β-sheet formation among _179_CVNITV_184_ fragments of sheep prion protein [17]. These observations align with the conventional role of organic solvents as denaturing and precipitating agents for proteins. Conversely, other studies showed that ethanol can inhibit the condensation of amyloid-β (Aβ) 1-42 and lysozyme [18, 19], effectively preventing amyloid dimerization and reducing toxicity of Aβ in cell lines[18]. In addition to biomolecular condensates, alcohols also have divergent effects on the formation of complex coacervates. Some studies reported that alcohols enhance the formation of coacervates [20–23], while others showed that alcohols weaken their formation [11, 24, 25]. These studies highlight our limited understanding of the interactions between condensates and solvents, as well as the need for further systematic investigation.

To resolve this conundrum, it is vital to understand the physicochemical specificities of condensates [15, 26]. For example, different condensates have distinct microenvironments that enable the selective compartmentalization and distribution of molecules within cells [27, 28]. The physicochemical specificities of condensates are fundamentally engendered by their sidechain composition and molecular interactions, such as electrostatic interactions between oppositely charged groups and cation-π, π-π interactions associated with aromatic groups [2, 29, 30]. Notably, recent studies have revealed that electrostatic interactions and other non-electrostatic interactions, including cation-π, π-π, and hydrophobic interactions, exhibit different responses to variations in temperature and salt concentration [7, 31]. For example, while electrostatic interactions can be screened by salts, non-electrostatic interactions can be significantly strengthened at a high salt concentration [7, 32, 33]. These studies highlight the unique roles of sidechain chemistry-encoded interactions in governing the responsiveness of condensates to environmental stimuli [31, 34].

In this study, we systematically investigate the dependence of phase transitions of condensates on their sidechain chemistry and molecular interactions. Using minimalistic coacervate models, in combination with systematic experiments and atomistic simulations, we demonstrate that hydrophilic coacervates primarily comprised of charged groups and electrostatic interactions are solidified by alcohols. In contrast, hydrophobic coacervates containing many aromatic groups and non-electrostatic interactions, such as cation-π and π-π interactions, are melted. These principles are further demonstrated to be applicable to biomolecular condensates formed by intrinsically disordered proteins and folded proteins, where the phase transition depends on the amino acid compositions. These findings highlight different nature of molecular interactions in dictating physicochemical microenvironments within condensates, offering promising insights for developing therapeutic drugs targeting pathological condensates and engineering stimuli-responsive biomaterials from the perspective of molecular interactions.

## Results and discussion

### Alcohol-induced phase transitions of condensates depend on sidechain chemistry

In response to alcohol molecules, condensates can undergo liquid-to-sold or solid-to-liquid phase transitions (LSPT and SLPT), as shown in Fig. 1a. In this work, we select alcohol molecules as a representative solvent stimuli and study how they modulate phase transitions of condensates (Fig. 1a). From a molecular perspective, the physical properties of condensates are the result of collective interactions between constituent polymers and solvents. The constituent polymers have different sidechains and interacting motifs with distinct hydrophobicity, and solvents can have different polarity (Fig. 1d). Considering different hydrophobicity of polymers, we hypothesize that condensates as a collective ensemble of polymers should also possess different degree of “hydrophobicity” depending on their sidechain chemistry and molecular interactions involved. Specifically, condensates predominantly composed of charged sidechains and electrostatic interactions tend to be hydrophilic (Fig. 1e), whereas condensates having abundant hydrophobic aromatic sidechains and cation-π, π-π interactions may be relatively hydrophobic (Fig. 1f).

**Figure 1.**
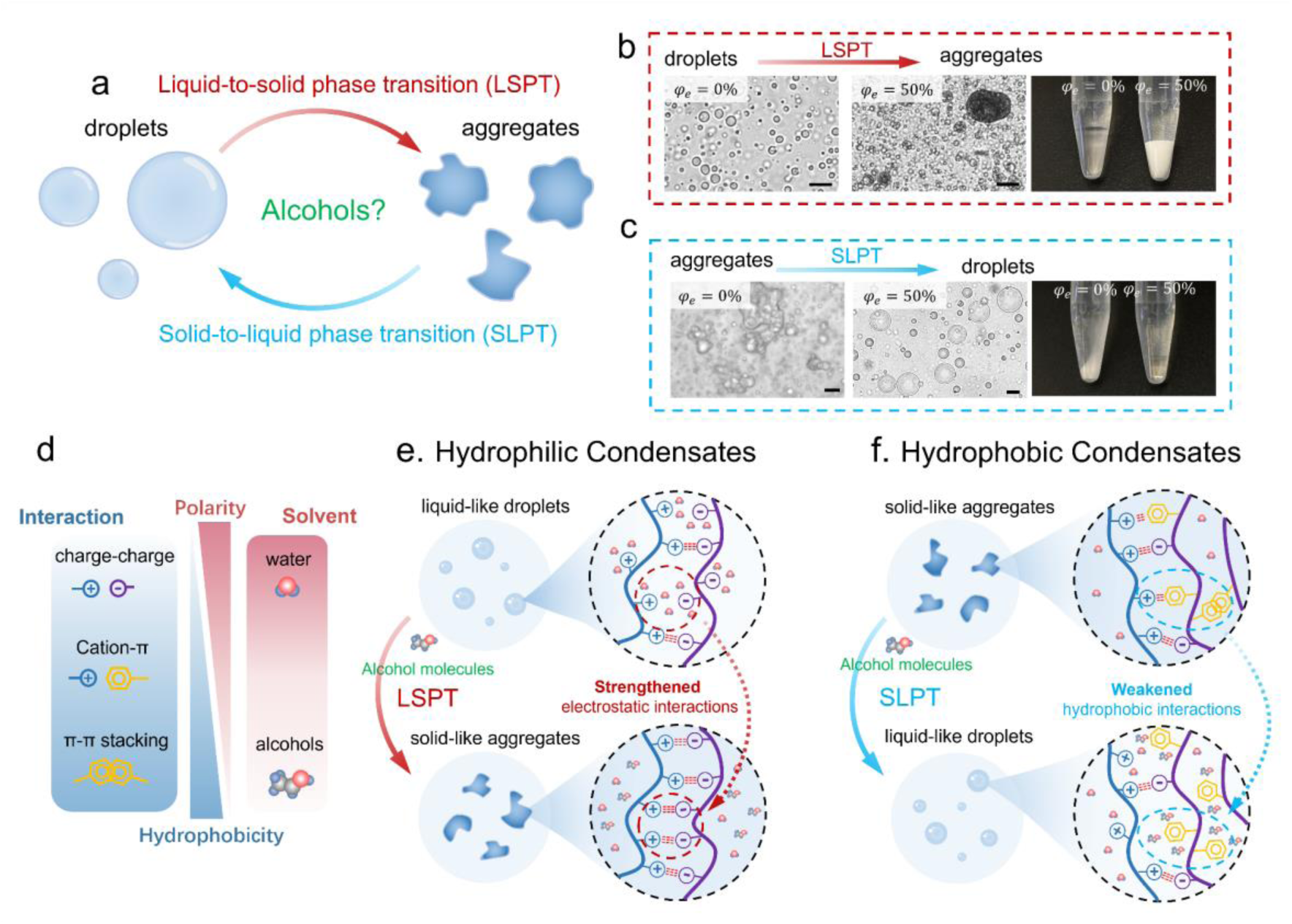
Sidechain chemistry-encoded biphasic phase transitions of condensates modulated by alcohols. (a) Schematic illustration of liquid-to-solid or solid-to-liquid phase transitions (LSPT or SLPT) of condensates modulated by alcohols. (b) Microscopic bright-field images and macroscopic photographs of LSPT of coacervates formed by negatively charged DEX-Sulf (2.5 wt%) and positively charged DEAE-DEX (5 wt%), as the volume fraction of ethanol increases from *φ_e_* = 0% to *φ_e_* = 50%. (c) Microscopic bright-field images and macroscopic photographs of SLPT of coacervates comprised of negatively charged PSS and positively charged PDDA with increasing *φ_e_* from = 0% to = 50% at the presence of 1.5 M NaCI. The concentration of charged monomer *c_m_* is 50 mM for PSS and PDDA. Scale bars in (b) and (c) are 20 μm. The chemical structures of these charged polymers in (b) and (c) are shown in Supplementary Fig. 1. (d) Schematic illustration of the relative polarity and hydrophobicity of different molecular interactions and solvents. (e) Hydrophilic condensates consist of mainly charged sidechains transition from liquid-like droplets to solid-like aggregates upon the introduction of alcohols. This liquid-to-solid phase transition (LSPT) is driven by strengthened electrostatic interactions. (f) Hydrophobic condensates having many aromatic groups are melted by alcohols, transitioning from solid-like aggregates into liquid-like droplets. This solid-to-liquid phase transitions (SLPT) is triggered due to weakened non-electrostatic interactions (e.g., cation-π and π-π interactions).

Theoretically, hydrophilic condensates could be solidified by alcohol molecules due to strengthened electrostatic interactions, leading to an LSPT. For example, the electrostatic free energy *f_el_* can be calculated using a generalized Debye−Hückel expression [35, 36], showing that *f_el_* scales with the dielectric constant solvents *ε_s_* as *f_el_*∼ −(*ε_s_*)−^3/2^ (See supplementary Note 1 for details). As a result, introducing alcohol molecules causes a decrease of *ε_s_*, leading to increased *f_el_* and thus strengthened electrostatic interactions. In contrast, hydrophobic condensates could be melted as a result of weakened hydrophobic interactions, resulting in SLPT. Apart from electrostatic interaction between two charges, all other interactions involve polarization effects [37]. These interactions are collectively contributed by van der Waals interactions arising from instantaneous fluctuations in electron distributions under the influence of an external electric field [37]. The energetic contribution from van der Waals interactions are approximated as 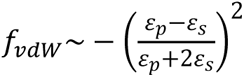 is the permittivity of polymers (See supplementary Note 1 for details). It is found that the absolute value of *f_vdW_* decreases with decreasing *ε_s_*, suggesting weakened van der Waals interactions. From another perspective, the hydrophobic interactions are weakened because the ethyl groups of alcohols could solubilize nonpolar hydrophobic compounds and interfere their interactions [38–40]. These theoretical arguments offer a hypothetical framework that the alcohol-induced phase transition of condensates depends on their sidechain chemistry and involved molecular interactions.

To support the hypothesis, we first select two representative coacervates with different sidechain chemistry and molecular interactions. We then add ethanol to trigger the phase transition of these coacervates at a volume fraction denoted as *φ_e_*=*V_e_*/(*V_e_* + *V_w_*), where *V_e_* and *V_w_* are volumes of ethanol and water, respectively. The first coacervate is formed by dextran derivatives, negatively charged Dextran sulfate sodium (DEX-Sulf) and positively charged Diethylaminoethyl-dextran (DEAE-DEX) (Fig. 1b). This coacervate is expected to be hydrophilic due to the presence of abundant hydrophilic hydroxyl groups (Supplementary Fig. 1). The bright-field image show that these coacervate behaves as liquid-like droplets in water, and the macroscopic dense phase is transparent settling at the bottom of a microtube after the centrifugation (Fig. 1b, *φ_e_* = 0%). However, introducing ethanol (*φ_e_* = 50%) into the solution leads to the formation of solid-like aggregates and LSPT (Fig. 1b and Supplementary Video 1), along with an opaque, dense phase. Rheology experiments further show that the storage modulus (*G*^′^) and loss modulus (*G*″) of DEX-Sulf/DEAE-DEX coacervates increase by more than three orders of magnitude in the presence of ethanol (Supplementary Fig. 2). Thermogravimetric analysis (TGA) reveals that this increase in moduli is due to the loss of solvent and the increased weight ratio of polyelectrolytes within the coacervates (Supplementary Fig. 3).

In comparison, another coacervate is formed by more hydrophobic polyelectrolytes that are positively charged polydiallyldimethylammonium chloride (PDDA) and negatively charged poly sodium 4-styrenesulfonate (PSS) containing aromatic groups (Supplementary Fig. 1). In water, PSS/PDDA coacervates exhibit as solid-like aggregates and the dense phase is relatively opaque (Fig. 1c, *φ_e_* = 0%). Upon introducing ethanol (*φ_e_* = 50%) into the solution, these coacervates transition into liquid-like droplets and the dense coacervate phase becomes transparent (Fig. 1c, *φ_e_* = 50% and Supplementary Video 2). Rheology experiments reveal that *G*^′^ and *G*^″^ of PSS/PDDA coacervates decreases significantly for more than 4 orders of magnitude upon ethanol introduction (Supplementary Fig. 2). Consistently, TGA shows that the volume fraction of solvents increases in coacervate phase after introducing ethanol, resulting in more swelled coacervate structures (Supplementary Fig. 2). Intriguingly, solid-like PSS/PDDA aggregates formed at *φ_e_* = 0% can bear heavy loads (Supplementary Fig. 4), whereas their counterparts formed at *φ_e_* = 50% readily drip off from a spoon, displaying typical liquid behaviors (Supplementary Fig. 4).

Moreover, other alcohol molecules, including 1,6 hexanediol, methanol, and isopropanol, can similarly cause LSPT and SLPT for the DEX-Sulf/DEAE-DEX coacervate and PSS/PDDA coacervate, respectively, suggesting a general underlying mechanism (Supplementary Fig. 2 and Supplementary Fig. 5). Collectively, these experiments support our hypothesis that condensates with different sidechains and molecular interactions exhibit completely opposite phase transitions in response to alcohol molecules.

### Aromatic sidechains underlie the hydrophobicity of coacervates and lead to solid-to-liquid phase transition

Aromatic groups have been shown to play critical roles in dictating the formation and physical properties of biomolecular condensates [30, 41–43] and complex coacervates [11, 44]. We further perform systematic experiments to validate the effect of aromatic sidechain on the hydrophobicity of coacervates and thus their phase transitions. We select two polyanions for comparison analysis, PSS and Poly (vinylsulfonic acid, sodium salt) (PVS), where only PSS monomer contains an aromatic group but monomers of PVS do not have (Fig. 2a). PDDA and poly-L-lysine (PLL) are chosen as polycations in complexation with PSS and PVS, respectively (Fig. 2a). Coacervates are all formed at a 1:1 neutral stoichiodmetry of charged monomers throughout the paper unless otherwise mentioned.

**Figure 2.**
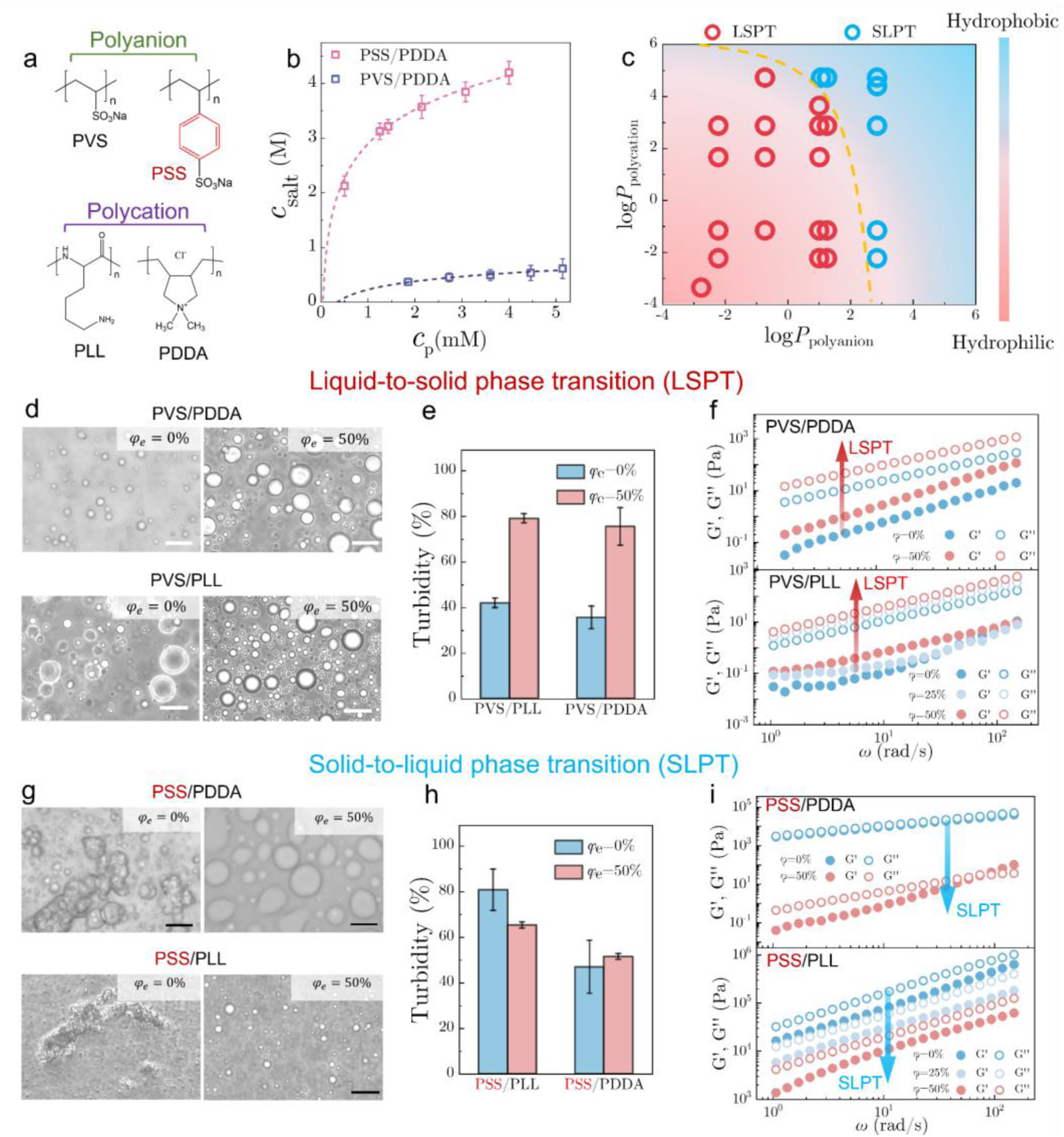
Aromatic residues encode the chemical specificity and phase transition of complex coacervates modulated by alcohols. (a) Chemical structures of two polyanions, PSS and PVS, and two polycations, PLL and PDDA. The aromatic sidechain of PSS is marked in red to highlight its structural difference compared to PVS. (b) Binodal curves of PSS/PDDA and PVS/PDDA complex coacervates showing the stability of coacervates against monovalent salts. *c_p_* and *c_salt_* are the molar monomer concentration for polymers and NaCl, respectively. The coacervation regime is below the binodal curve. Sodium chloride (NaCI) is used as the salt. (c) Plot of partition coefficient (*logP*) of polyanions and polycations of coacervates in the current work and literature. Values of *logP* are predicted by a cheminformatics-based software [45] by inputting the monomeric structure of polymers. A yellow dash lines demarcate the regime of SLPT and LSPT. (d) Bright-field images of PVS/PDDA (*c_m_* = 50 *mM* and *c_s_* = 0 M) and PVS/PLL (*c_m_* = 50 *mM* and *c_s_* = 0 M) coacervates at *φ_e_* = 0% and *φ_e_* = 50%. (e) Turbidity of PVS/PDDA and PVS/PLL coacervate solutions at *φ_e_* = 0% and *φ_e_* = 50%, respectively. (f) Rheology frequency sweeps of f PVS/PDDA and PVS/PLL coacervates at different *φ_e_*. Samples are measured with a constant strain γ=0.5 % within the linear viscoelastic regime. (g) Bright-field images of PSS/PDDA (*c_m_* = 50 MM and *c_s_* = 1.5 M) and PSS/PLL (*c_m_* = 50 *mM* and *c_s_* = 0 M) coacervates at *φ_e_* = 0% and *φ_e_* = 50%. (h) Turbidity of PSS/PDDA and PSS/PLL coacervate solutions at *φ_e_* = 0% and *φ_e_* = 50%, respectively. (i) Rheology frequency sweeps of f PVS/PDDA and PVS/PLL coacervates at different *φ_e_*. Samples are measured with a constant strain γ=0.5 % within the linear viscoelastic regime. The scale bars in (c) and (f) are 20 μm.

The binodal curves, below which phase separation occurs, are firstly determined using sodium chloride (NaCI) for PSS/PDDA and PVS/PDDA coacervates. We find that the two-phase area of PSS/PDDA coacervates is much broader than that of PVS/PDDA coacervates (Fig. 2b). A similar trend is consistently observed when measuring the binodal curve using another potassium chloride (KCI) salt (Supplementary Fig. 6). The stability of PSS/PDDA coacervates against salts is contributed by additional aromatic residue-associated cation-π and π–π interactions. Besides their stability, they also adapt different physical morphologies; PVS/PDDA coacervates behave as liquid-like droplets in water (*φ_e_* = 0%) (Fig. 2d), while PSS/PDDA coacervates exhibit as solid-like aggregates (Fig. 2g). Substituting PDDA with PLL yields analogous outcomes, with PVS/PLL shown as liquid-like droplets (Fig. 2d) but PSS/PLL behaved as solid-like aggregates (Fig. 2g).

In line with our hypothesis, introducing ethanol (*φ_e_* = 50%) causes the transition of solid-like PSS/PDDA and PSS/PLL aggregates into liquid-like droplets (Fig. 2g). Conversely, PVS/PDDA and PVS/PLL coacervates show negligible changes in their morphologies after introducing ethanol (Fig. 2d). However, it is noted that after introducing ethanol PVS/PDDA and PVS/PLL coacervate droplets are less likely to wet solid substrate, evident by decreased wetting areas on untreated glass slides (Supplementary Fig. 7). In addition, the solution turbidity of these two coacervates is observed to increase after the introduction of ethanol (Fig. 2e); however, the turbidity of PSS/PDDA and PSS/PLL coacervates slightly decreases (Fig. 2h). These results validate that the phase separation of PVS/PDDA and PVS/PLL coacervates is enhanced by ethanol.

To quantitatively evaluate impact of ethanol on physical properties of coacervates, rheology experiments are performed. It is firstly found that the storage modulus *G*^′^ and loss modulus *G*^″^ of PSS/PDDA and PSS/PLL coacervates are several orders of magnitude higher than those of PVS/PDDA and PVS/PLL coacervates (Fig. 2f and Fig. 2i), consistent with their different microscopic physical morphologies (Fig. 2d and Fig. 2g). In particular, *G*^′^ and *G*^″^ of PSS/PDDA and PSS/PLL coacervate are nearly equal (Fig. 2i), indicative of their physical properties resembling solid-like gels. After introducing ethanol at different volume fractions *φ_e_*, *G*^′^ and *G*^″^ of PSS/PDDA and PSS/PLL coacervates significantly decrease by more than four and two orders of magnitude, respectively (Fig. 2i). Conversely, the addition of ethanol leads to an increase in *G*^′^ and *G*^″^ of PVS/PDDA and PVS/PLL coacervates by approximately two orders of magnitude (Fig. 2f). These experiments using comparative coacervate systems and various characterization techniques consolidate that aromatic sidechains encode distinct chemical specificities of coacervates, thereby inducing different phase transitions by alcohols.

To reinforce the claim, we substitute PDDA with another polycation, poly[(vinylbenzyl) trimethylammonium chloride] (PVBTMA), which shares similar chemical structures to PDDA but includes an aromatic residue. Consistently, it is evident that *G*^′^ and *G*^″^of PVS/PVBTMA coacervates decrease upon the introduction of ethanol (Supplementary Fig. 8a), in contrast with the increased *G*^′^ and *G*^″^ of PVS/PDDA lacking aromatic residues (Fig. 2f). We have explored many other coacervates without aromatic residues, including poly (N,N-dimethylaminoethyl methacrylate) (PDMAEMA) and poly(acrylic acid) (PAA), DEX-Sulf and PLL, Adenosine triphosphate (ATP) and PLL. Remarkably, consistent with our hypothesis, all of these systems undergo LSPT after introducing alcohols, as characterized by rheology or optical microscopy (Supplementary Fig. 9).

Furthermore, we map out the hydrophobicity of polycations and polyanions into a phase separation diagram based on all previously reported and our current coacervate systems (Fig. 2c). The hydrophobicity of these polymers is represented by their partition coefficient (*logP*) in a typical octanol-water system and predicted by a cheminformatics-based algorithm [45] by inputting the monomeric structure of polymers. It is noted that we focus on evaluating the hydrophobicity of sidechains and thus we designedly block charged groups during the calculation of *logP*. The higher the *logP*, the more hydrophobic the substance is. Notably, this phase diagram reconciles literature results showing that the regimes of LSPT and SLPT can be well distinguished based on the hydrophobicity of polymers (Fig. 2c). These results also consolidate the importance of sidechain hydrophobicity in indicating chemical specificities and thus the responsiveness of condensates.

### Time-ethanol superposition reveals changes in interaction strength

To better understand the variations in physical properties and molecular interactions within coacervates affected by alcohols, we employ a superposition method to analyze the rheology data. Superposition is a well-known rheological principle to study the response of viscoelastic materials over a broad range of frequencies [20, 46]. We first conduct frequency sweeps on two exemplary complex coacervates, PAA/PDMAEMA and PVS/PVBTMA, at different volume fractions of ethanol and temperatures. The overall shape of rheology curves of these coacervates is typical of polymer melt with viscoelastic properties being dominated by the loss modulus (*G*^″^) at low frequency and the storage modulus (*G*^′^) at high frequency (Supplementary Fig. 8 and Supplementary Fig. 10).

Consistent with the previous observations and our hypothesis, the presence of ethanol leads to significant decrease in *G*^′^and *G*^″^ of PVS/PVBTMA coacervates containing aromatic residues, whereas it increase *G*^′^and *G*^″^ of PAA/PDMAEMA coacervates lacking aromatic residues (Supplementary Fig. 8). By employing a time-ethanol superposition where the frequency and modulus are multiplied by a critical relaxation time (*τ_c_*) and a vertical shift factor *b_φ_*, frequency sweeps conducted at different volume fractions of ethanol (*φ_e_*) are rescaled onto master curves (Fig. 3a and Fig. 3b). The successful application of time-alcohol superpositions implies that these coacervates remain self-similar and homogeneous structures. Therefore, studying the viscoelastic properties of coacervates at different conditions is equivalent to examining them at different time scales [20, 47]. Intriguingly, in the time-ethanol superpositions, *τ_c_* and *b_φ_* of PVS/PVBTMA coacervates decrease, while those of PAA/PDMAEMA coacervates increase. In particular, the trend of *τ_c_* over *φ_e_* can be effectively captured by linear functions in semilog plots (Fig. 3d and Fig. 3e), in line with previous studies [20]. The *τ_c_* is physically determined by an energy barrier *E* of associated interactions between stickers on polymer chains and environmental temperature *T*, as *τ_c_*∼exp (*E*/*k*_B_*T*), where *k*_B_ is the Boltzmann constant [46]. With this relationship, these results suggests that interaction strength within coacervates scale linearly with the volume fraction of ethanol. Differently, the interaction strength within PAA/PDMAEMA coacervates increase over *φ_e_*, while that within PVS/PVBTMA coacervates decreases.

**Figure 3.**
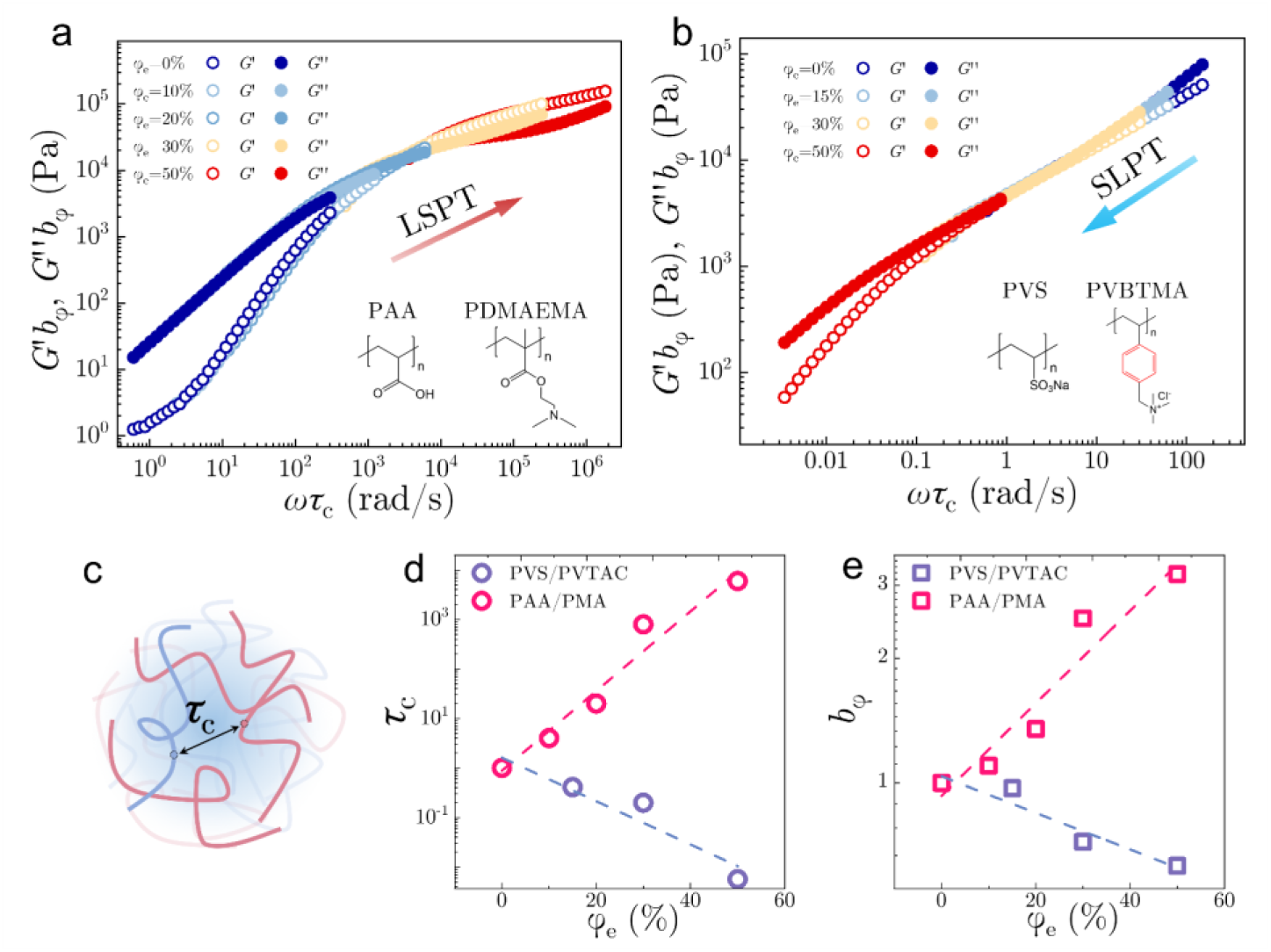
Time-alcohol superposition. Time-alcohol superposition plots of (a) PAA/PDMAEMA and (b) PVS/PVBTMA complex coacervates. The insert images show the chemical structures of polyelectrolytes. The concentration of charged monomer *c_m_* is 50 mM for all polyelectrolytes and coacervates are therefore formed at a neutral stoichiometry. Frequency sweeps conducted at various volume fraction of ethanol (*φ_e_*) are used for the time-alcohol superposition (Supplementary Fig. 8). The LSPT and SLPT are identified for these two coacervates, respectively. (c) Schematic illustration of the critical relaxation time *τ_c_* which is related with the interactions between associative residues. (d) (e) Values of the critical relaxation time *τ_c_* and vertical shift factors *b_φ_* as a function of *φ_e_*. The dashed lines are linear fittings to experimental points in the semi-logarithmic plots.

Moreover, it is found that *G*^′^ and *G*^″^ of these two coacervates monotonically decrease with increasing temperature (Supplementary Fig. 10). In addition, the critical frequency at which *G*^′^ equals *G*^″^ shift toward higher frequencies, suggesting faster relaxation and more liquid-like properties at the high temperature (Supplementary Fig. 10). By applying a time-temperature superposition, frequency sweeps at different temperatures can be rescaled onto a single master curve using *T* = 20 °C as the reference point (Supplementary Fig. 11), with 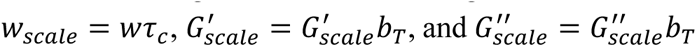. It is observed that *τ_c_* and *b_T_* continually decrease with increasing temperature (Supplementary Fig. 11). The trend of *τ_c_* over *T* can be well fitted by the relationship of *τ_c_*∼exp (*E*/*k*_B_*T*) in a semilog coordinate (Supplementary Fig. 11). These results suggest that temperature can induce a melting of coacervates, regardless of their different sidechain compositions.

### Isothermal titration calorimetry reveals variations in thermodynamic parameters of coacervate formation

We further apply isothermal titration calorimetry (ITC) to understand the influence of alcohols on the driving forces responsible for the formation of coacervates. In ITC experiments, a high-concentration titrant within a syringe is gradually loaded, binding to another analyte in a sample cell (Fig. 4a). Heat flows required to maintain a constant temperature during the complexation process are monitored to extract the change of thermodynamic parameters during the reaction. ITC experiments are conducted for the comparative coacervate systems in Fig. 2, using the polycation as the analyte and the polyanion as the titrant.

**Figure 4.**
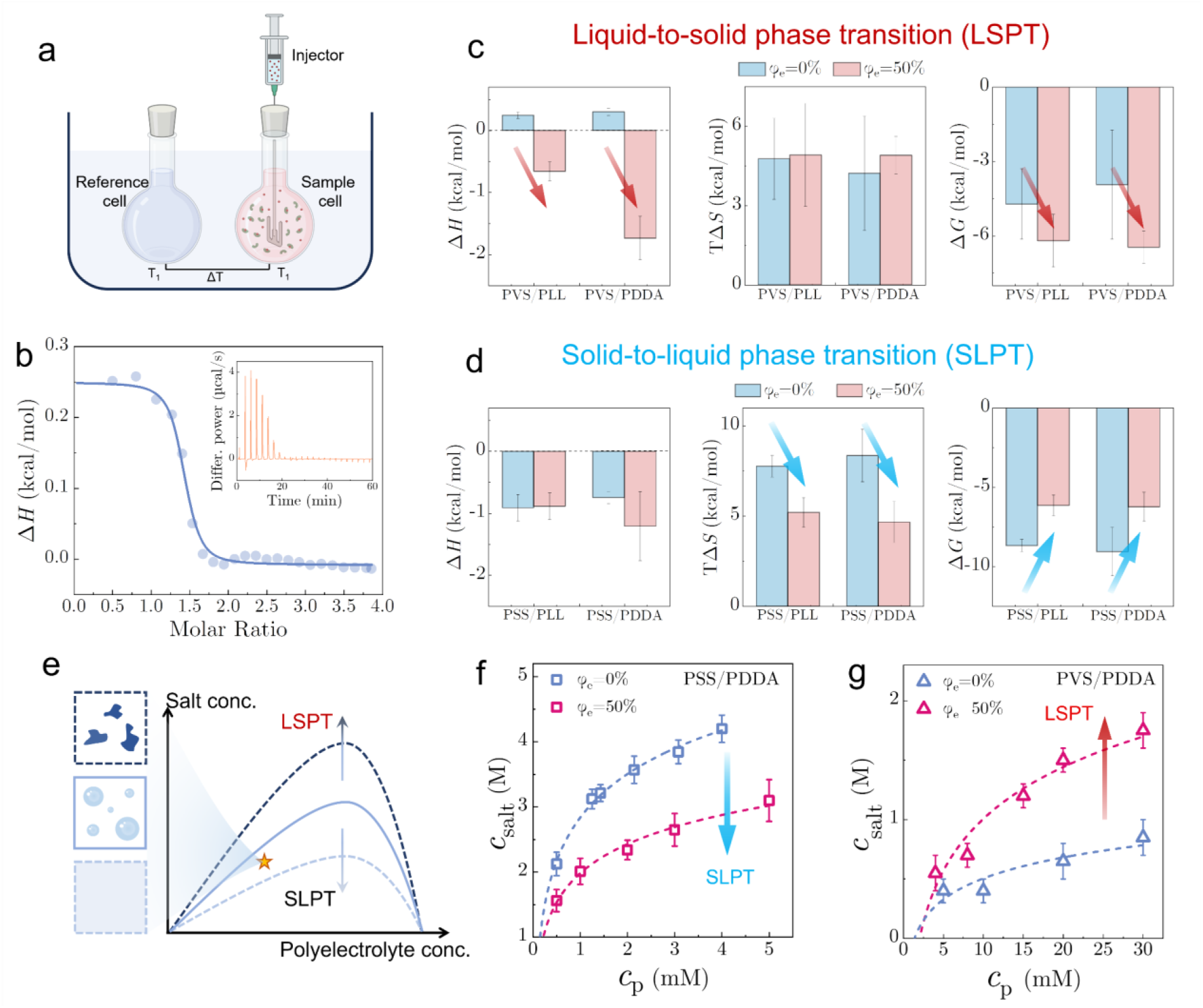
Microcalorimetry reveals changes of entropy, enthalpy, and Gibbs free energy. (a) Schematic illustration of configurations of isothermal titration calorimetry. (b)The molar enthalpy change Δ*H* versus time for the isothermal injection of PVS (*c_m_* = 1 MM) into a cell containing PLL (*c_m_* = 20 MM). The blue circles are experimental data, which are fitted using a one-site binding model. The insert image shows the differential power (DP) signal versus time. (c) Molar enthalpy Δ*H*, entropy times temperature *T*Δ*S*, and Gibbs free energy Δ*G* for PVS/PLL and PVS/PDDA coacervates which undergo LSPT. (d) Molar enthalpy Δ*H*, entropy times temperature *T*Δ*S*, and Gibbs free energy Δ*G* for PSS/PLL and PSS/PDDA coacervates which undergo SLPT. (e) Schematic illustration showing that different phase transitions lead to different shifting of the binodal curves and variation in the physical morphologies of coacervates. Experimental phase diagram of (f) PSS/PDDA and (g) PVS/PDDA coacervates at *φ_e_* = 0% and *φ_e_* = 50%. Sodium chloride is used as the salt. Error bars in (f) and (g) indicate mean ± SD (n = 3 independent samples). From the phase diagram, PSS/PDDA coacervates exhibit a SLPT, while PVS/PDDA coacervates show a LSPT.

The raw thermographs, in general, exhibit typical exothermic curves for PSS/PDDA and PSS/PLL coacervates containing aromatic residues, irrespective of the presence of ethanol (Supplementary Fig. 12). Each exothermic peak corresponds to a single injection of titrant. However, in the absence of ethanol, several endothermic peaks are identified, followed by a plateau approaching zero. This phenomenon is attributed to the formation of solid aggregates resulting from strong interactions between macromolecules [48, 49], as previously observed in Fig. 2c. In comparison, the complexation of PVS/PDDA and PVS/PLL coacervates without aromatic residues is characterized as entropy-driven endothermic processes (Fig. 4b and Supplementary Fig. 13). This endothermic process is triggered by the release of hydration water from dissolved polyelectrolytes chains during the complexation [50]. However, in presence of ethanol (*φ_e_* = 50%), the complexation process of PVS/PDDA and PVS/PLL become exothermic (Supplementary Fig. 13), suggesting that electrostatic interactions are enhanced.

The enthalpy change for each injection is determined by integrating and normalizing the peaks in thermographs with respect to time and the amount of material injected. By fitting the enthalpy results using a one-site binding model, we can quantitatively extract thermodynamic parameters, including changes in enthalpy (Δ*H*), entropy (Δ*S*), and Gibbs free energy (Δ*G*) (Fig. 4c and Fig. 4d). Results reveal that Δ*H* of PSS/PDDA and PSS/PLL coacervates containing aromatic residues is not significantly affected by the addition of ethanol (Fig. 4d). However, the changes of entropy (Δ*S*) and Gibbs free energy (Δ*G*) of these coacervates decreases (Fig. 4d). This decreased entropy could be caused by the solvation of hydrophobic groups by ethanol molecules [38–40]. Conversely, ethanol has minimal impact on the entropy of coacervates lacking aromatic residues, such as PVS/PDDA and PVS/PLL coacervates (Fig. 4c). However, the introduction of ethanol causes the enthalpic change (Δ*H*) for these two coacervates to transition from positive to negative (Fig. 4c), indicating strengthened molecular interactions. Also, the change of Gibbs free energy (Δ*G*) for these coacervates increases (Fig. 4c). Collectively, the ITC results suggest that the SLPT is primarily caused by the entropy change in the formation of coacervates, whereas the LSPT is an enthalpy-driven process due to more favorable electrostatic interactions.

The change of Gibbs free energy is correlated with the phase diagram [51, 52]. In the case of complex coacervates, the phase diagram is often delineated as a plot of polyelectrolyte concentration against salt concentration (Fig. 4e). The single-phase and two-phase regimes are demarcated by a binodal curve, below which complex coacervation occurs. To further validate the effect of ethanol on phase separation, we measure the phase diagram of PSS/PDDA and PVS/PDDA in the absence and presence of ethanol. Generally, a large phase separation area is indicative of a stronger tendency to phase separate and thus a greater reduction in Δ*G* [52, 53]. Indeed, the results show that the phase separation area of PSS/PDDA coacervates decreases in the presence of ethanol (Fig. 4f), consistent with the observed diminished Δ*G* by ITC (Fig. 4d). In contrast, the phase separation area of PVS/PDDA coacervates expands (Fig. 4g), aligning with the increment of Δ*G* after the introduction of ethanol (Fig. 4c).

### Atomistic simulations reveal the changes in molecular interactions

We further employ molecular dynamics simulations to understand how interactions are modulated by ethanol within coacervates having different sidechain compositions. In simulations, polyelectrolytes are represented by their monomers for simplicity (Fig. 5b). Monomers of PSS and PVS are selected due to their minimal difference, as PSS contain one additional benzene ring structure compared to PVS. The monomer of PLL is constructed to interact with monomers of PSS and PVS. 20 pairs of oppositely charged monomers are solvated by purely water molecules or a mixture of water and ethanol molecules in a simulation box with the periodic boundary condition (Fig. 5a). 20 pairs of monovalent sodium chloride salts are inserted to ensure the electroneutrality of the system. Details of simulation setups are described in he Materials and Methods section.

**Figure 5.**
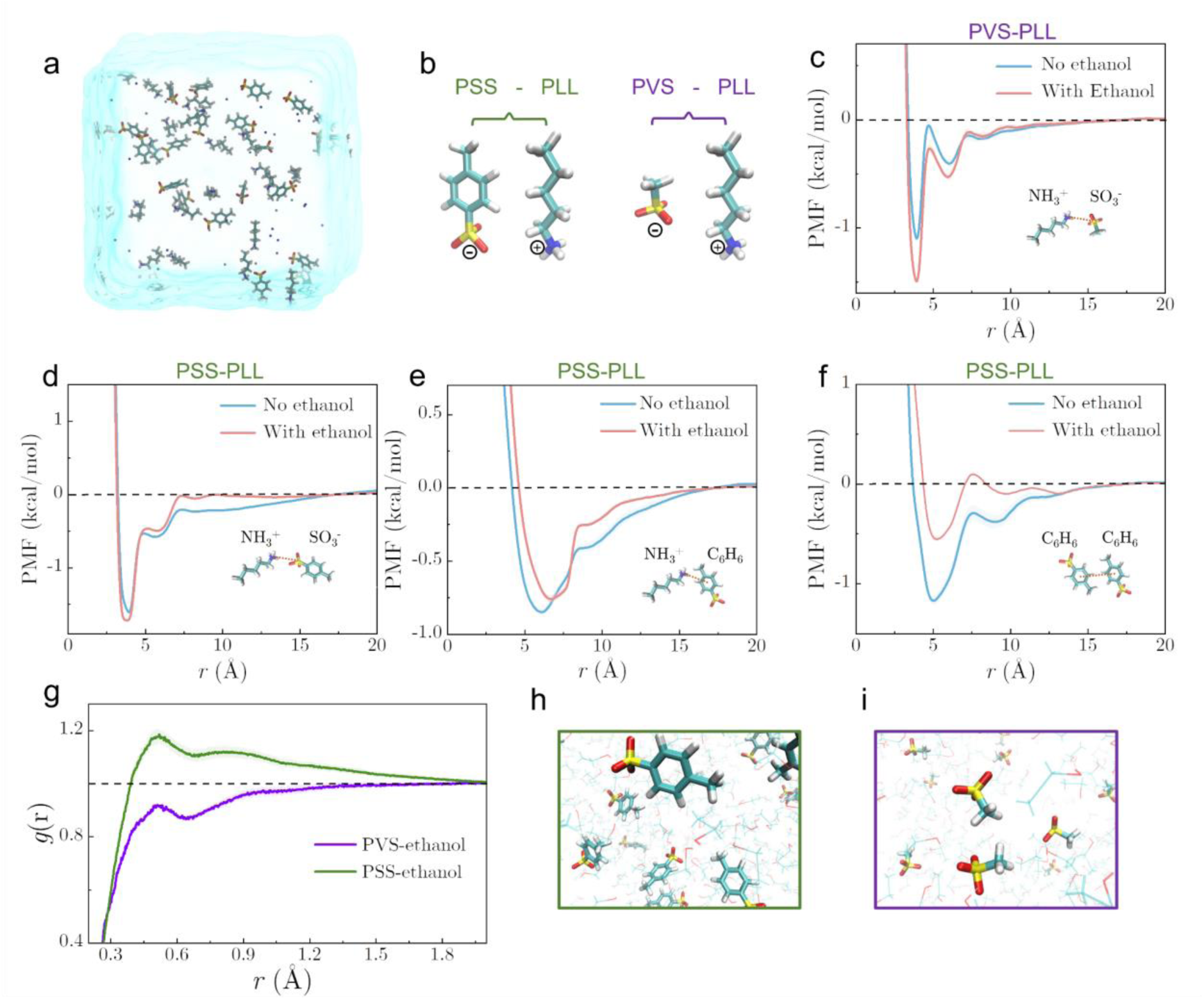
All-atom molecular dynamic simulation of complex coacervates. (a) Schematic illustration of the simulation configuration. The box contains oppositely charged monomers, salts, and water/ethanol. (b) Chemical structures of charged monomers of PSS, PVS, and PLL. The potential of mean force (PMF) between selected charged monomers in explicit solvent conditions as a function of center-of-mass distance for (c) PVS/PLL and (d) (e) (f) PSS/PLL systems. The insert images in (d) (e) (f) show the selected coordinates for calculating PMF. (g) Radial distribution function of ethanol molecules around PVS and PSS. Snapshots shows the distribution of ethanol molecules around (h) PSS and (i) PVS molecules. For all curves, standard deviations (shaded areas) are obtained for all simulations across 5 independent replicates with different random seeds.

Fromm these simulations, we extract the potential of mean forces (PMF) as a function of the distance between different interacting sidechains of monomers. The results show that PMF between oppositely charged NH_3_^+^ of PVS and SO_3_^-^ of PLL increases in the presence of ethanol (Fig. 5c). In addition, we confirm that the dielectric constant surrounding media decrease after incorporating ethanol molecules (Supplementary Fig. 14). Conversely, the attraction between NH_3_^+^ of PSS with a benzene ring structure and SO_3_^-^ of PLL is slightly weakened by the introduction of ethanol (Fig. 5d). Although a minimal decrease is observed for the deepest potential well occurring at a small molecular distance, the absolute value of integration of PMF over the interaction distance is decreased. Moreover, by choosing different reaction coordinates, it is found that the PMFs of NH_3_^+^-C_6_H_6_ and C_6_H_6_-C_6_H_6_ decrease clearly in the presence of ethanol (Fig. 6e and Fig. 6f). These results demonstrate that the electrostatic interaction is strengthened while other interactions, such as cation-π and π-π, are weakened by ethanol molecules.

**Figure 6.**
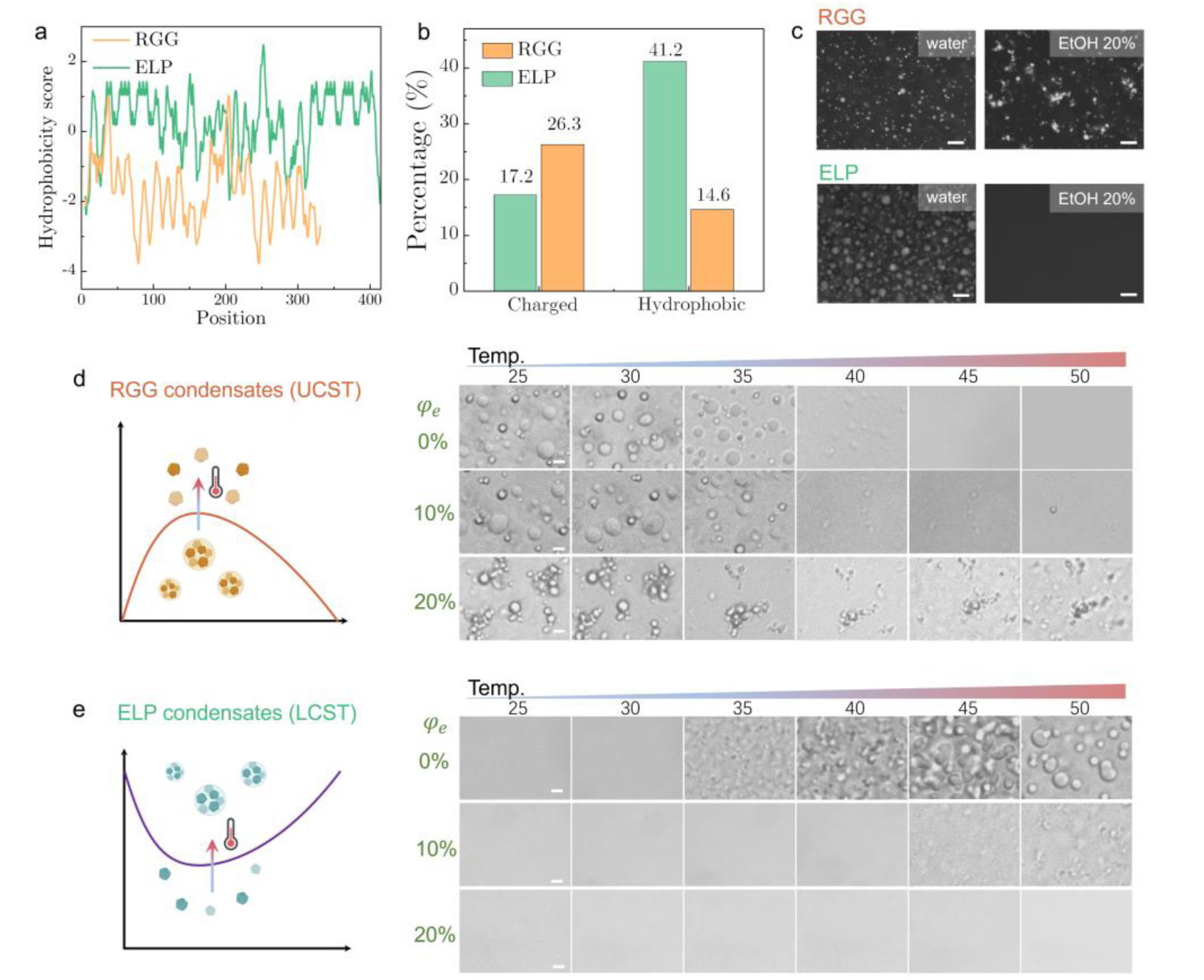
Amino acid composition-dependent phase transitions of condensates. (a) The hydrophobicity score of amino acids for the RGG and ELP proteins using a widely used Kyte-Doolittle scale [54]. (b) The percentage of charged and hydrophobic amino acids for the RGG and ELP proteins. (c) Fluorescence images of condensates formed by the RGG (15 µM) and ELP proteins (67 µM) in pure water and water/ethanol mixture (*φ_e_* = 20%). The RGG protein is connected to a green GFP protein and the ELP protein is labeled with Cy3 dye. Condensates are formed in 0.1 M PBS solution with 150 mM NaCI. The scale bars are 20 µm. (d) Schematics of RGG condensates that are dissolved when the solution temperature is above the upper critical solution temperature (UCST). Bright-field images of RGG condensates at different *φ_e_* of ethanol and temperatures. (e) Schematics of RGG condensates that are dissolved when the solution temperature is above the upper critical solution temperature (UCST). Bright-field images of ELP condensates at different *φ_e_* of ethanol and temperatures. The scale bars in (d) and (e) are 5 µm.

In addition, ITC experiments also suggest the weakening of coacervates containing aromatic residues has large entropic contributions. As explained, this entropic contribution is likely caused by the redistribution and solvation of ethanol around hydrophobic groups [38–40]. To demonstrate this, we measure the radial distribution function (RDF) of ethanol with respect to PSS and PVS by simulations, denoted at *g*(*r*). RDF is typically defined as the ratio of the probability of finding a particle at a certain distance to the probability of finding a particle in a homogeneous system. When *g*(*r*) > 1, it indicates an excess probability of finding molecules compared to a random distribution. Conversely, *g*(*r*) being less than 1 means that the likelihood of finding particles at that distance is lower than expected in a homogenous system. RDF results show that ethanol molecules are more likely to distribute around and solvate PSS that has a benzene ring structure compared to PVS. In particular, at distances larger than 0.4 nm, *g*(*r*) of PSS-ethanol is above 1 (Fig. 5g). In contrast, *g*(*r*) of PVS-ethanol is constantly below 1 at all distances (Fig. 5g). This implies that ethanol is favourable in dispersing closely around PSS. In addition, it is found that ethanol can better solvate PSS than water (Supplementary Fig. 15a). These results explain the large entropic variation in PSS/PLL coacervates modulated by ethanol (Fig. 4d). However, the minimal difference is identified between the solvation of PVS by ethanol and water (Supplementary Fig. 15b). These results support that the complexation between PVS and PLL is primarily driven by the enthalpic electrostatic interaction rather than changes in solvation energy, as shown in Fig. 4c. Overall, these atomistic simulations provide molecular-level explanations that qualitatively agree with experimental results, consolidating our hypothesis that coacervates having different sidechain chemistry and molecular interactions exhibit different alcohol-induced phase transitions.

### Amino acid composition-dependent phase transitions of biomolecular condensates

Next, we extend our hypothesis to biomolecular condensates of intrinsically disordered proteins (IDPs) and show that their phase transitions, modulated by ethanol, depend on amino acid composition. Two IDPs, namely RGG and elastin-like polypeptide (ELP), are expressed and purified. The RGG protein is derived from LAF-1 proteins which is the main component of membrane-less P granules. The ELP protein is composed of repeating amino acid sequences that mimic the structure of elastin, a protein found in connective tissue that provide elasticity and resilience.

We first estimate the hydrophobicity of IDPs using a widely used Kyte-Doolittle scale, which defines the relative hydrophobicity of amino acid residues [54]. The more positive the value, the more hydrophobic the amino acids are. Consequently, it is found that the ELP protein has higher average hydrophobicity scores compared to the RGG protein (Fig. 6a). In addition, compared to the ELP protein, the RGG protein has more charged amino acids, such as arginine and aspartic acid, and fewer hydrophobic amino acids, including valine and leucine (Fig. 6b). These results suggest that the RGG protein is more hydrophilic, while the ELP protein is more hydrophobic.

Next, we form condensates using these two IDPs and study how ethanol affects their morphologies and thermo-responsiveness. First, the RGG protein form liquid-like droplet condensates at room temperature (Fig. 6c). These RGG condensates dissolve when the solution temperature is above an upper critical solution temperature (UCST) (Fig. 6d). However, introducing ethanol at only 20% volume fraction (*φ_e_* = 20%) cause a solidification of RGG condensates, suggesting a LSPT. In contrast, the ELP protein does not phase-separate at room temperature but forms liquid-like droplet condensates when the temperature is above a lower critical solution temperature (LCST), which is around 35 °C in our experiments. Differently, the introduction of ethanol (20%) leads to a complete dissolution of ELP condensates, exhibiting an SLPT. Besides affecting their physical morphologies, ethanol also tunes the critical temperature, UCST or LCST, for these two types of condensates. Specifically, RGG condensates exhibit higher UCST as introducing more ethanol and condensates becoming solid-like aggregates. ELPs condensates also have an increasing LCST, but condensates can not form anymore with increasing volume fraction of ethanol.

Additionally, ethanol has caused biphasic phase transitions for protein-polymer condensates. Two widely used folded proteins, bovine serum albumin (BSA) and lysozyme, are selected to interact with polyelectrolytes, PDDA and PAA, respectively. They both form liquid-like droplet condensates in water at a neutral PH (Supplementary Fig. 16). It is observed that BSA/PDDA condensates become solid-like aggregates with increasing volume fraction (*φ_e_*) of ethanol (Supplementary Fig. 16a). In addition, the solution turbidity continues to increase over *φ_e_* (Supplementary Fig. 16b). These results suggest that ethanol cause a LSPT for BSA/PDDA condensates. In contrast, lysozyme/PAA condensates are completely dissolved by ethanol at an intermediate volume fraction *φ_e_* = 25%, followed by a re-entrant formation of solid-like aggregates at *φ_e_* = 50% (Supplementary Fig. 16c). This re-entrant phenomenon is also observed in the solution turbidity over *φ_e_* (Supplementary Fig. 16d). Interestingly, this re-entrant phase separation is also identified for coacervates consist of aromatic residues, such as PSS/PDDA and PVS/PVBTMA (Supplementary Fig. 17). These results are consistent with our analysis and previous studies showing that the BSA protein is more hydrophobic than the lysozyme protein (Supplementary Fig. 18) [55, 56]. In addition, another hydrophobic Amyloid β Protein Fragment 1-42 (Aβ 1-42), which are widely identified in the pathogenesis of Alzheimer’s disease[57, 58], is also examined. In aqueous solutions (*φ_e_* = 0%), Aβ 1-42 form solid-like plaque structures in water labeled by Thioflavin T (ThT) (Supplementary Fig. 19). Interestingly, these solid-like Aβ 1-42 plaques are completely dissolved within 30 seconds upon introducing ethanol at *φ_e_* = 50% (Supplementary Fig. 19). Overall, these results demonstrate that the phase transition of biomolecular condensates modulated by ethanol closely depends on the amino acid composition of proteins.

## Conclusion

The present work combines systematic experimental characterizations and atomistic simulations to demonstrate that the chemical specificities of condensates depend on their sidechain compositions. As a result, the distinct chemical specificities of condensates lead to different phase transitions, namely LSPT or SLPT, in response to the introduction of alcohol molecules. Importantly, the framework of sidechain chemistry-dependent bidirectional phase transition resolves the puzzle in the literature regarding the inconsistent effects of alcohols on condensates or coacervates. In addition, our work suggests that condensates containing hydrophobic residues exhibit internal environments reminiscent of organic solvents, which agrees with previous studies [59]. In contrast, condensates primarily comprised of hydrophilic residues encompass internal environments in close proximity to water. These different chemical environments may be crucial for condensates to selectively sequester small molecules and dynamically modulate enzymatic reactions [6, 28]. Moreover, the chemical specificities of condensates are also correlated with their physical morphologies, such as liquid-like droplets or solid-like aggregates. For example, solid-like aggregates are often melted by alcohol molecules into liquid-like droplets, suggesting their proximate chemical nature [59]. These insights provide new understandings of the physicochemical properties of condensates and offer promising strategies for designing therapeutic small-molecule drugs that can reverse or mitigate solid-like aggregates based on the sidechain composition of constituent components [15].

Our work also reveals that the phase transition of condensates is fundamentally governed by the interplay of different types of molecular interactions. For instance, the electrostatic interactions are enhanced by alcohol molecules, while non-electrostatic interactions, such as cation-π and π-π interactions, can be weakened. This finding echoes with the previous work showing that the electrostatic interaction and non-electrostatic interactions exhibit different responses to monovalent salts and temperature [7, 31]. These results highlight the distinct nature of electrostatic interactions versus non-electrostatic interactions; however, the underlying mechanisms causing these differences remain to be investigated in future studies. The new understanding that these two types of interactions show different responses to environmental stimuli is instrumental for designing functional condensates or other materials with stimuli-responsive properties, such as those with LCST or UCST, for various industrial and biomedical applications [34].

## Materials and Methods

### Materials

Poly(sodium 4-styrenesulfonate) (PSS, Sigma-Aldrich, 243051); Poly(diallyldimethylammonium chloride) (PDDA, Sigma-Aldrich, 522376); Poly-(L-lysine) (PLL, Macklin, P885950); Poly(vinylsulfonic acid, sodium salt) (PVS, Macklin, P909334); Poly(methacrylic acid, sodium salt)(PMA, Sigma-Aldrich, 434507); Poly(vinylbenzyl trimethylammonium chloride) (PVBTMA, Scientific Polymer Products, 707); Poly(N,N-dimethylaminoethyl methacrylate) (PDMAEMA, Polymer Source, P16059); Poly(acrylic acid) (PAA, Polymer Source, P14708); Ethanol (xxxx), methanol (Sigma-Aldrich, 34860), isopropanol (Sigma-Aldrich, PX1838P); Sodium chloride 5M, RNase-free (ThermoFisher, AM9760G); KCl 2 M, RNase-free (ThermoFisher, AM9640G); Bovine Serum Albumin (BSA, Sigma-Aldrich, SRE0096); Amyloid *β* Protein Fragment 1-42 (amyloid-β 1-42, Macklin, A834109); Lysozyme (Macklin, L6051); DEAE-Dextran (Macklin, D873343); Dextran sulfate sodium salt (DEX-sulf, Macklin, D808272); hexadecane (Macklin, H854563); Millipore Milli-Q water (18.2 MΩ, pH=7) is used in all experiments. Sodium chloride, sodium dihydrogen phosphate, potassium chloride, disodium phosphate, nickel sulfate and imidazole were brought form Macklin. SDS was brought from Bio-RAD. The ultra-pure glycerol was brought from Invitrogen. LB broth, TB broth and agarose were purchased from Thermo Scientific Fisher. Plasmid extraction was performed using Takara Plasmid Miniprep Kit or E.Z.N.A.® Plasmid Mini Kit I, (V-spin) from Omega Bio-tek company. PCR and restriction enzymes digest were carried out by Takara EmeraldAmp® GT PCR Master Mix and Thermo FastDigest enzymes. DNA fragments were purified by gel electrophoresis using Bio-Rad. DNA Electrophoresis Cells & Systems and extracted through TaKaRa MiniBEST Agarose Gel DNA Extraction Kit or Omega E.Z.N.A.® Gel Extraction Kit. Fragments ligation was achieved by Thermo T4 DNA ligase or Takara T4 DNA ligase. FPLC instrument (GE Healthcare, New York, US) installed with a HisTrap HP His tag protein purification columns (Cytiva, US). The sonication of the protein was carried out in ultrasonic crusher (Wuxi Voshin instruments manufacturing Co., LTD). All materials were used as received without further purification.

### Coacervate formation

Stock solutions of each polyelectrolyte were prepared initially at high concentrations using Milli-Q water (18.2 MΩ, pH=7). Complex coacervates were prepared by mixing oppositely charged polyelectrolytes at a neutral 1:1 stoichiometry ratio of monomer charge between polycations and polyanions. The mixing solutions were vigorously vortexed for approximately 20 seconds. The turbid solutions were centrifuged at 4000 g for 10 minutes to separate the coacervate phase from the dilute phase. The coacervate phases were extracted for further rheology and TGA characterizations.

### Optical microscopy

The optical images of complex coacervates were captured by an inverted fluorescence microscope (Olympus). The coacervate solutions with a volume of around 8 μl were loaded into chambers that were customized using a spacer (ThermoFisher, S24737) sandwiched by two coverslides at a 120 μm gap. The coverslides were first cleaned by ethanol and water to remove contaminants.

### Phase diagram

The left arm of the binodal curve was measured by titrating the salt solution of high concentration into the coacervate solution prepared at different initial polymer concentrations. The polyelectrolyte and salt concentrations at the critical point where the coacervate solution became transparent as identified by naked eyes were recorded. For each critical point on the binodal curve, at least three independent experiments were conducted to minimize errors.

### Turbidity measurements

The turbidity of solutions was measured in 96-well plates using a SpectraMax iD3 Multi-Mode Microplate Reader (Molecular Devices, CA). After mixing oppositely charged polyelectrolytes, the solution was gently stirred using a pipette and quickly transferred into a 96-well plate with a volume of 200 μl. The turbidity was calculated by normalizing the absorption values of coacervate solutions by their maximum values at λ=580 nm.

### Rheology experiment

The rheology experiments were performed using a stress-controlled Anton Paar MCR301 rheometer. Coacervates samples were loaded onto the bottom surface of rheometer and a cone plate with a diameter of 25 mm and a cone angle of 1° was used as the top geometry. The top plate was lowered onto samples keeping a fixed gap of 0.051 mm. The excess materials were trimmed off. Hexadecane was used to seal samples and the plate to reduce evaporation of water. The temperature was controlled at 20 °C using a Peltier temperature control system coupled to the bottom surface of the rheometer. Frequency sweeps of coacervates were conducted with the applied strain at between 0.1 % and 1%, which was confirmed to be within the linear viscoelastic regime by strain sweep experiments.

### Thermogravimetric Analysis

Approximately 15-30 mg of coacervate samples were loaded into crucibles whose weights were recorded beforehand. After loading the samples, the weights of the crucibles were measured again using a micro-balance to determine the weight of the coacervates. The experiments were performed using a TGA analyzer (DZ TGA101 Dazhan). The furnace was first heated up to 70 °C at a rate of 20 °C/min and kept at 70 °C for 10 minutes to evaporate all ethanol. Afterwards, the temperature was increased to 110 °C at a rate of 20 °C/min for 20 minutes to evaporate water. The weights of the crucibles were recorded over time and used to calculate the weight ratios of ethanol, water, and polyelectrolytes with ions, respectively. For each type of coacervate, at least three independent samples were prepared and measured to calculate the mean value and standard deviation.

### Fourier transform infrared spectroscopy

FTIR was performed using a PerkinElmer Spectrum Two spectrophotometer with a LiTaO3 detector. Measurements were performed in Attenuated total reflectance (ATR) mode after a calibration with a clean background. The spectra were collected at wave number ranging from 600 to 4000 cm^-1^ at 4 cm^-1^ resolution averaging 20 scans per sample. Data analysis was conducted using the Spectrum 10 software.

### Isothermal titration calorimetry

Calorimetric measurements were performed using a MicroCal PEAQ-ITC Automated (Malvern) instrument with a sample cell volume of 200 μL. An injection syringe of 40 μL was used to load polyelectrolytes at a concentration that was typically 15-25 times higher than the concentration of oppositely charged polyelectrolyte in the sample cell. As the standard control, equivalent DI water was titrated into the reference cell to clean the system at least twice before starting formal experiments. For each experiment, the temperature is set to 25 °C, and a constant stirring rate of 750 rpm was applied on a paddle-shaped tip of the injection syringe. The adsorption heat was calculated by subtracting the baseline. In all cases, solvent compositions in syringe and sample cell were identical to minimize mixing or dilution heat. ITC data was fitted with the one-site binding model using MicroCal PEAQ-ITC Analysis software (Version 1.0.0.1259). For each reaction, trial experiments were conducted to determine the optimal experimental conditions.

### All-atom molecular dynamic (MD) simulations

All simulations were performed using the GROMACS 2021 package [60, 61] on the GPU nodes of HPC2021 at The university of Hong Kong. The chemical structures of small molecules (PSS, PVS, PLL, and ethanol) were built using the Avogadro software [62]. The force field parameters and topologies of small molecules were derived by the ACPYPE program [63] using amber2 for the “atom type” option. The partial charges of each molecule were fit based on the quantum mechanics calculation using the RESP program [64] on an online R.E.D. Server [65]. Data analysis and image export were performed using the VMD software [66].

For each simulation, 20 pairs of oppositely charged small molecules were solvated in a cubic water box with the size of 5.2×5.2×5.2 *nm*^3^ and periodic boundary conditions. Then 20 Na^+^ and 20 CI^-^ ions were added to the systems. For simulations with ethanol, 700 ethanol molecules were inserted into the box to replace water molecules. The number ratio of ethanol to water molecules is around 1:3. The TIP3P water model was used [67]. The force field parameters for Na^+^ and CI^-^ ions were adapted from the reference [68]. The overall energy of the system was minimized using the steepest descent method followed by 500 ps NVT and 500 ps NPT equilibration simulations at 300 K and 1 bar. The equilibrated structure was used as the starting structure for MD simulation at 300 K and 1 bar. The simulation time step is 2 fs and the total simulation time is 500 ns. The V-rescale thermostat [69] was used for maintaining the temperature with a coupling time constant of 0.1 ps. The Parrinello-Rahman method [70, 71] was used to maintain the pressure at 1 bar with a coupling time constant of 2.0 ps. PME algorithm [72, 73] was applied to calculate the long-rang electrostatic interactions. The cutoffs of electrostatic potential and van der Waals potential were set to 1.2 nm. The LINCS algorithm was used to constrain all bonds with H-atoms [74]. MATLAB scripts were developed to analyze the radius distribution function *g*(*r*) and calculate PMF as −*kTlog*(*g*(*r*)). Five independent simulations were performed for each condition with random initial positions.

### Plasmid construction

Two plasmids used in this study are listed in the supplementary Table 2. E. coli TOP10 was used for cloning and plasmid propagation and grown in selective Luria-Bertani medium or Luria Bertani plates with 1.5 wt% agar. Antibiotics of 100 µg/mL ampicillin were added for selection. The DNA sequences of the plasmid constructs containing PCR-fragments were confirmed by sequencing.

### Expression and purification of protein

Recombinant plasmids were transformed into E. coli BL21 (DE3) for protein expression. Seed culture was prepared by picking up the single colony into 3 mL LB with corresponding antibiotics and shaked at 37 °C overnight. Then, 1% (v/v) overnight culture was added into sterilized Terrific broth (TB) medium at 37 °C with 220 rpm until OD reached 0.6-0.8 with antibiotics. Isopropyl β-D-1thiogalactopyranoside (IPTG) was added to induce the protein expression. After that, cultures were consecutively incubated at 17 °C and 180 rpm for overnight protein expression. The cells were harvested by centrifugation (4000 g for 10 min) and the pellets were washed twice with buffer and then frozen at -80 °C for the following purification. Proteins including RGGRGGEGFP and SpyTagELPCarHELP SpyTag were bonded with HiTrap His column for further purification. SDS-PAGE was applied to test the expression and purification of proteins, and the protein content was calculated with a BCA kit in which BSA is as the standard protein.

For the purification of proteins, the cell pellets were suspended in lysis buffer (0.1 M PBS buffer, pH 7.4, containing 1 M NaCl, 20 mM imidazol) and lysed with ultrasonic treatment for 30 min in an ice bath. The supernatant after centrifuging at 12, 000 g about 30 min was filtered with 0.22 µm filter membrane, and was bind with nickel his column for 1 h. The purification was conducted with FPLC, the protein was eluted with elution buffer (0.1 M PBS buffer, pH 7.4, containing 1 M NaCl, 500 mM imidazol) at a step gradient. The eluted proteins were collected and concentrated with Millipore of 10 kDa, and then, the proteins was buffer changed with storage buffer (0.1 M PBS buffer, pH 7.4, 150 mM NaCl) and was freezed with liquid nitrogen for storage at -80 °C.

### Labeling of proteins

For Cy3 labeling, the protein spytag-elp-carH-elp-spytag was dissolved in 0.1 M sodium bicarbonate buffer, pH 8.3 at a concentration of 2 mg/mL. NHS-Cy3 ester dissolved in dimethyl sulfoxide was slowly added with stirring. Protein was added to Cy3 at the molar ratio of 1:1, and the mixture was incubated overnight at 0 °C. The Cy3 labeled proteins were then separated on the Zeba™ Spin Desalting Columns and exchanged to dissolve in PBS. The protein content was calculated with a BCA kit in which BSA is as the standard protein.

## Supporting information

Supplementary information for main text

## Acknowledgements

This work was supported by the General Research Fund (Nos. 17306221, 17317322, and 17306820) from the Research Grants Council (RGC) of Hong Kong, as well as the National Natural Science Foundation of China (NSFC)-RGC Joint Research Scheme (N_HKU718/19). H.C.S. was funded in part by the Croucher Senior Research Fellowship from Croucher Foundation and the Health@InnoHK program of the Innovation and Technology Commission of the Hong Kong SAR Government. X.Z. acknowledges the funding from the Research Grants Council of Hong Kong (No. 22302823) and National Natural Science Foundation of China (Young Scientist Fund No. 22303073). The MD computations were performed using research computing facilities offered by Information Technology Services, the University of Hong Kong.

## Author contributions

F.C. conceived the idea for this study. F.C., H.C.S. and X.Z. designed the study; F.C. performed the experiments and analyzed data with assistance from X.L., W.G. and C.W.; F.C. and X.Z. performed all-atom molecular dynamics simulations and analyzed the results. Y.H. and J.X. expressed and purified intrinsically disordered proteins; H.C.S. and X.Z. supervised the research. F.C. wrote the paper and all authors contributed to the revision of the paper.

