## Supplementary information for main text for "Sidechain chemistry-encoded solid/liquid phase transitions of condensates"

**Supplementary Table 1: Inconsistent effects of organic solvents on the phase transitions of coacervates or condensates reported in literature.**

| Material | Organic Solvent | Effect | reference |
|---|---|---|---|
| amyloid-β (Aβ) | Ethanol | SLPT | [1, 2] |
| Lysozyme | Ethanol | SLPT | [3, 4] |
| PSS/PVBTMA | Ethanol, ethylene glycol | SLPT | [5] |
| PAAcat/ QCS-Tf2N | DMSO | SLPT | [6] |
| PSS/P3BImHTBr | Tetrahydrofuran | SLPT | [7] |
| keratin-8 protein | Ethanol | LSPT | [8] |
| $_{179}$CVNITV$_{184}$ protein | Ethanol | LSPT | [9] |
| Chitosan/ Hyaluronic Acid | Ethanol, methanol | LSPT | [10] |
| PDDPC/PVS; PDDPC/PAA | Ethanol | LSPT | [11] |
| BPEI/PAA | Ethanol, methanol, and etc. | LSPT | [12] |

PSS: Poly(sodium 4-styrenesulfonate)

PVBTMA: Poly[(vinylbenzyl)trimethylammonium chloride]

PAAcat: Poly(acrylic acid) functionalized with catechols

QCS-Tf2N: Chitosan ion-paired with bis(trifluoromethane-sulphonyl)imide

P3BImHTBr: poly[3-[6´-(N-butylimidazolium)-hexyl]thiophene]bromide

PDDPC: Poly(N,N-dimethyl-3,5-dimethylene piperidinium chloride)

PVA: Polyvinylamine

PAA: Poly(acrylic acid)

PVS: poly(vinylsulfonic acid, sodium salt)

BPEI: branched poly(ethylene imine)

**Supplementary Table 2:** Plasmids and vectors used for expressing intrinsically disordered proteins.

| Plasmids | Vector |
|---|---|
| RGGRGG-EGFP | Pet 21 a |
| SpyTag-Elp-CarH-Elp-SpyTag | pQE-80L |

### Supplementary Note 1: Estimation of different driving forces affected by the introduction of organic solvents.

Molecular interactions, such as electrostatic interaction, cation - π, and π - π, are the main driving forces for the complex coacervation. Therefore, to investigate the effects of solvents on complex coacervations, we further examine the changes of molecular interaction energy based on theoretical models upon the introduction of organic solvents. For coacervates formed by oppositely charged polyelectrolytes with relatively hydrophilic sidechains, the main driving force for the phase separation is assumed to be the electrostatic interaction. For disconnected ions with diameters $\sigma = \nu^{1/3}$, electrostatic free energy is given by a generalized Debye−Hückel expression as[13, 14],

$$f_{el} = -\frac{1}{4\pi\nu}\left[\ln(1+\kappa\sigma) - \kappa\sigma + \frac{(\kappa\sigma)^2}{2}\right] \qquad (1)$$

where $\kappa$ is the inverse Debye screening length[15] given by $\kappa\sigma = \sqrt{4\pi(l_B/\sigma)\sum\phi_i f_i}$ with Bjerrum length $l_B = e^2/4\pi k_B T\varepsilon_0\varepsilon_s$ (where $f_i$ is the charge density of polymers or salts, $e$ is the elementary charge, $\varepsilon_0$ is the vacuum permittivity, $\varepsilon_s$ is the dielectric constant of the solvent). In the dilute limit, $\phi_i \rightarrow 0$, Eq. 1 reduces to well-known Debye−Hückel correlation free energy as used in Voorn-Overbeek (VO) theory[16],

$$f_{el} = -\kappa^3/12\pi. \qquad (2)$$

For simplicity, Eq. 2 is used to calculate electrostatic free energy in our work. Based on Eq. 2, $f_{el}$ scales with $(l_B)^{3/2}$ and thus $(\varepsilon_r)^{-3/2}$, as $f_{el} \propto (l_B)^{3/2} \propto (\varepsilon_r)^{-3/2}$. This indicates the electrostatic interaction is enhanced when the the dielectric constant of solvent decease upon the introduction of organic solvents into water.

For coacervates formed by oppositely charged polyelectrolytes with aromatic residues, cation-π and π - π interactions play important roles in the complex coacervation[17, 18]. Apart from electrostatic interactions between two charges, all other interactions involve polarization effects for molecules of polarizability $\alpha$. This is because all atoms and molecules are polarizable and the polarizability arise from the displacement of its negatively charged electron cloud relative to the positively charged nucleus under the influence of an external electric field [19]. Therefore, the energetic contributions from these short-range interactions in a solvent medium can be modeled as [19],

$$f_{sr} = -\frac{3kT}{r^6}\left(\frac{\varepsilon_p - \varepsilon_s}{\varepsilon_p + 2\varepsilon_s}\right)^2 \alpha^6, \tag{4}$$

where $\varepsilon_p$ and $\varepsilon_s$ are the dielectric consant of moleucles and solvents, respectively. Based on the Eq. 4, it is shown that $f_{sr}$ scales with $\left(\frac{\varepsilon_p - \varepsilon_s}{\varepsilon_p + 2\varepsilon_s}\right)^2$. The dielectric constant of water $\varepsilon_w$ and ethanol $\varepsilon_e$ at room temperature ($T = 20\ ℃$) are 80.2 and 24.3, respectively. The dielectric constant of a mixture of water and ethanol can be calculated as $\varepsilon_s = \varphi_e \varepsilon_e + (1 - \varphi_e)\varepsilon_w$ [20]. With this, $\varepsilon_s$ decrease monotonically with increasing $\varphi_e$ upon the introduction of ethanol into water. Therefore, it is found that $f_{sr}$ continues to decrease with decreasing value of $\varepsilon_s$, suggesting that the strength of cation-π and π - π interactions are reduced in the presence of organic solvents.

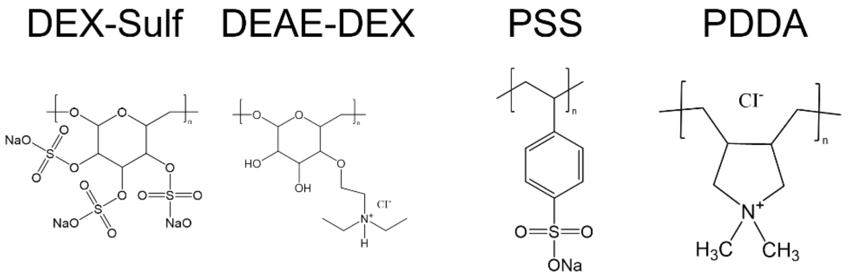


**Supplementary Fig. 1.** Chemical structures of DEX-sulf, DEAE-DEX, PSS, and PDDA polyelectrolytes.

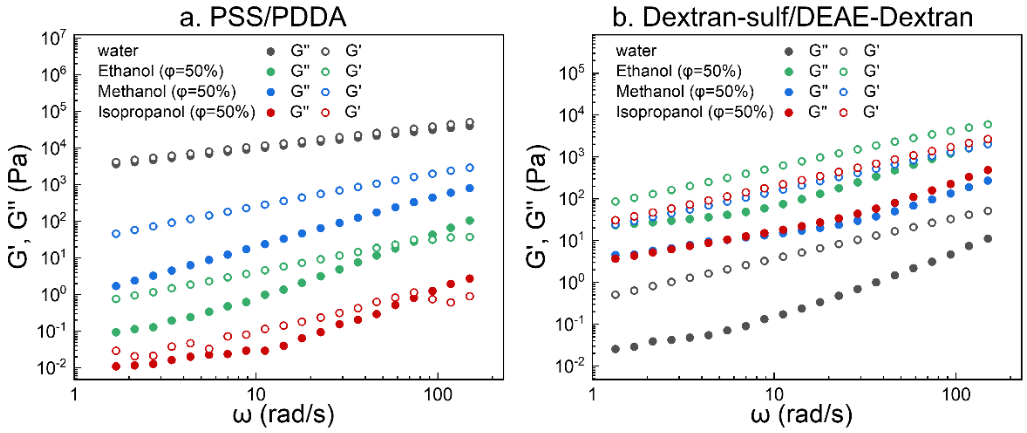


**Supplementary Fig. 2.** Frequency sweeps of (a) PSS/PDDA (100 mM, 1:1 stoichiometry of monomeric charge) and (b) Dextran-sulf (2.5 wt%)/DEAE-Dextran (5 wt%), formed at four different solvent conditions: water, water/ethanol (1:1 v/v), water/methanol (1:1 v/v), and water/isopropanol (1:1 v/v).

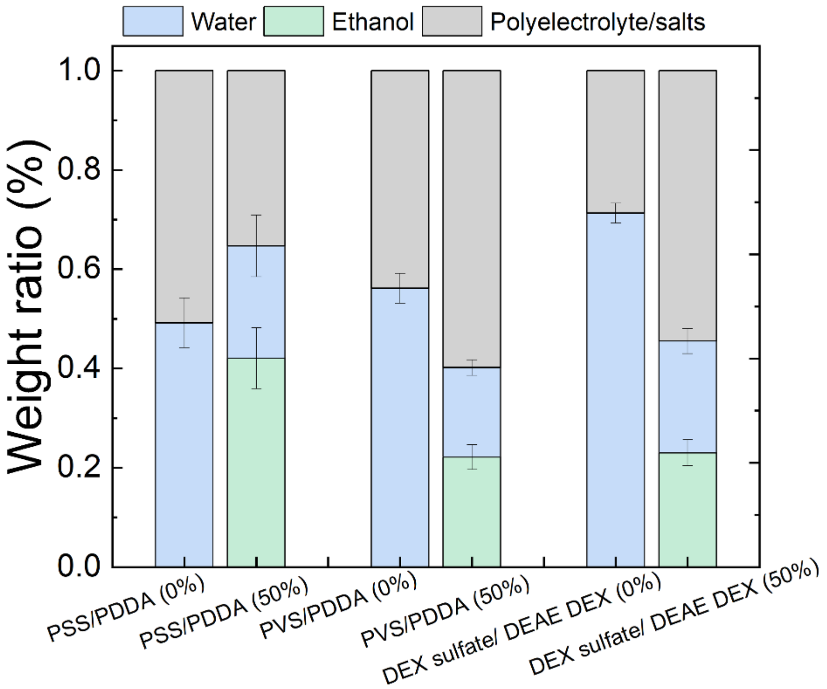


**Supplementary Fig. 3.** Weight ratios of water, ethanol, and polyelectrolyte/salt in different types of complex coacervates at aqueous solution and water/ethanol (1:1 v/v) mixture, measured by thermogravimetric Analysis. At least three independent samples are measured to calculate a mean value and standard deviation (error bars).

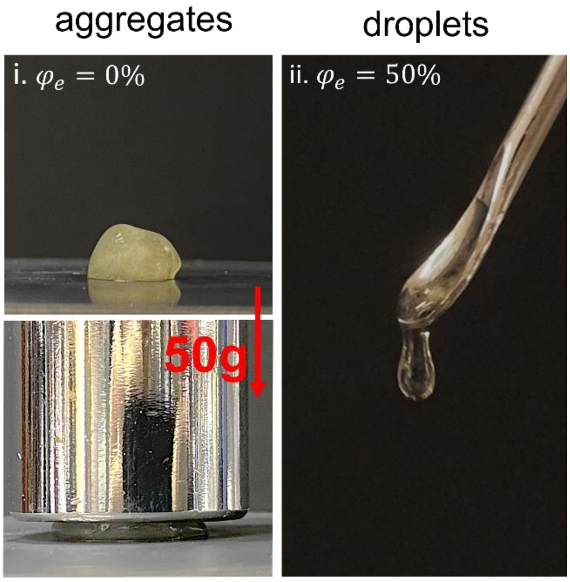


**Supplementary Fig. 4.** Photographs of PSS/PDDA coacervates formed at different volume fraction of ethanol. Coacervates exhibit as solid-like aggregates in pure water ($\varphi_e = 0\%$) and can bear heavy loads, while coacervates in the $\varphi_e = 50\%$ ethanol/water mixture behave as liquid which readily drip off from a spoon.

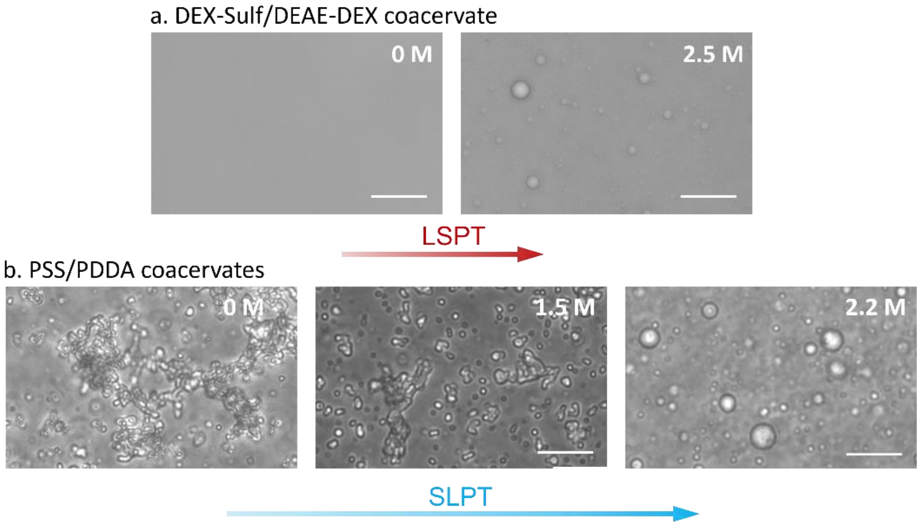


**Supplementary Fig. 5.** The effect of 1.6 hexanediol (HD) on coacervates with different sidechain compositions. (a) A liquid-to-solid phase transition (LSPT) is observed for DEX-Sulf (0.5 wt%)/DEAE-DEX (0.5wt %) coacervates. Coacervates do not form in pure water while they appears when the concentration of 1,6 HD is 2.5 M. (b) A solid-to-liquid phase transition (SLPT) for PSS/PDDA coacervates. The concentration of charged monomer is 5 mM for PSS and PDDA, and the concentration of salts (sodium chloride) is 500 mM. PSS/PDDA gradually transition from solid-like aggregates at 0 M 1,6 HD to liquid-like droplets at 2.2 M 1,6 HD.

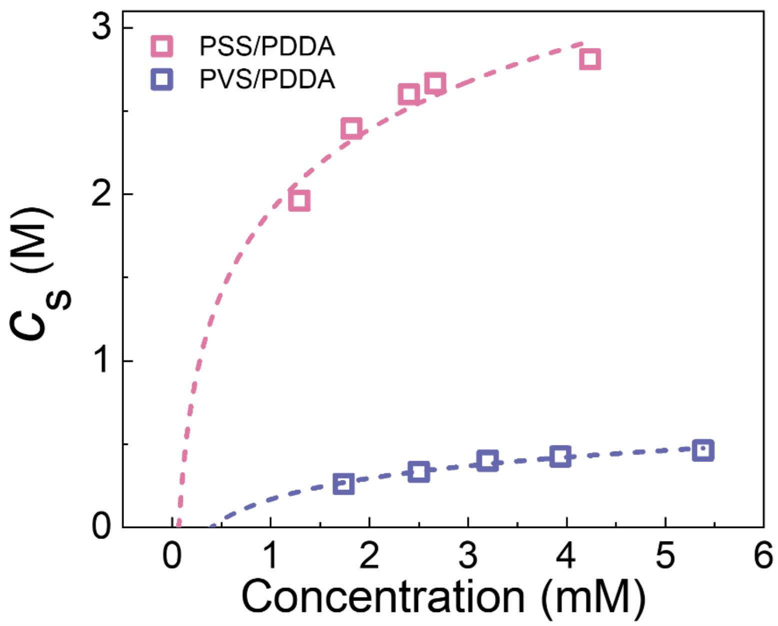


**Supplementary Fig. 6.** Left arms of phase diagram for PSS/PDDA (pink dash line) and PVS/PDDA (violet dash line) coacervates using the KCl salt. Coacervates are formed at a neutral stoichiometry ratio.

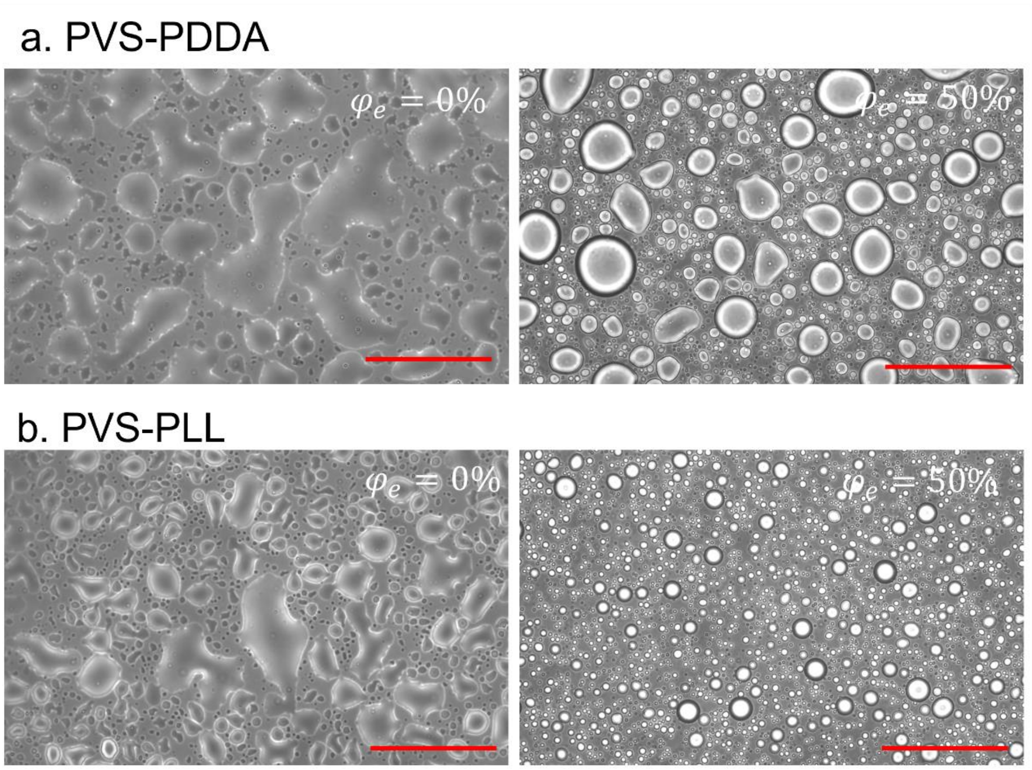


**Supplementary Fig. 7.** Bright-field images of (a) PVS/PDDA and (b) PVS/PLL coacervates. These images are recorded after pipetting solutions on cover slides for ~10 minutes, allowing the complete sedimentation and wetting of coacervate droplets. Scale bars are 100 μm.

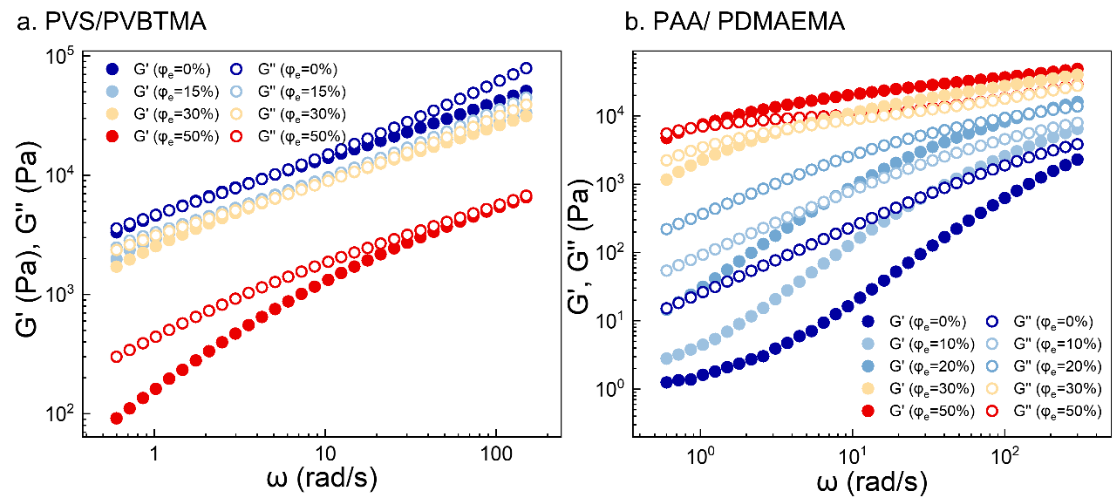


**Supplementary Fig. 8.** Frequency sweeps of (a) PVS/PVBTMA and (b) PAA/PDMAEMA coacervates at different volume fraction of ethanol ($\varphi_e$). The storage modulus $G'$ and loss modulus $G''$ of PVS/PVBTMA coacervates monotonically decrease over $\varphi_e$, while that of PAA/PDMAEMA coacervate increase as $\varphi_e$ increases.

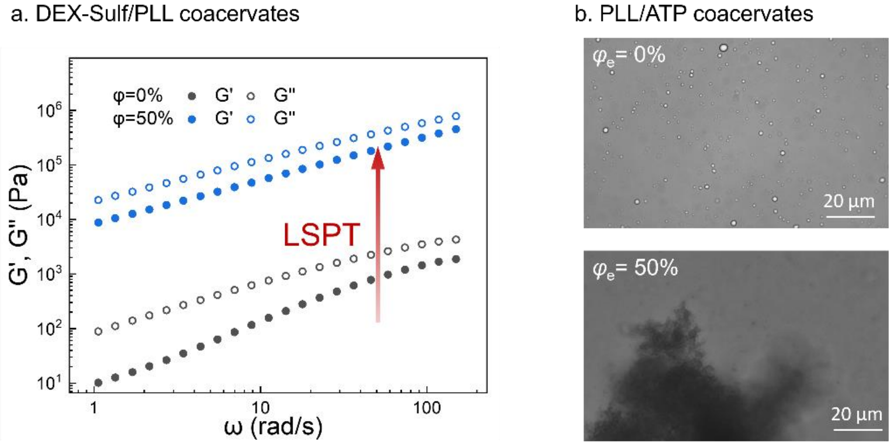


**Supplementary Fig. 9.** Experiments showing the liquid-to-solid phase transition of DEX-sulf/PLL and PLL/ATP coacervates. (a) Frequency sweeps of coacervates formed by DEX-sulf (2 wt%) and PLL (2 wt%) in water ($\varphi_e = 0\%$) and water/ethanol (1:1 v/v, $\varphi_e = 50\%$). The storage modulus $G'$ and loss modulus $G''$ increase by more than 2 orders of magnitudes after the introduction of ethanol. (b) Bright-filed images of coacervates formed by PLL and ATP in water ($\varphi_e = 0\%$) and water/ethanol (1:1 v/v, $\varphi_e = 50\%$). A phase transition from liquid-like droplets ($\varphi_e = 0\%$) to solid-like precipitates (1:1 v/v, $\varphi_e = 50\%$) is observed.

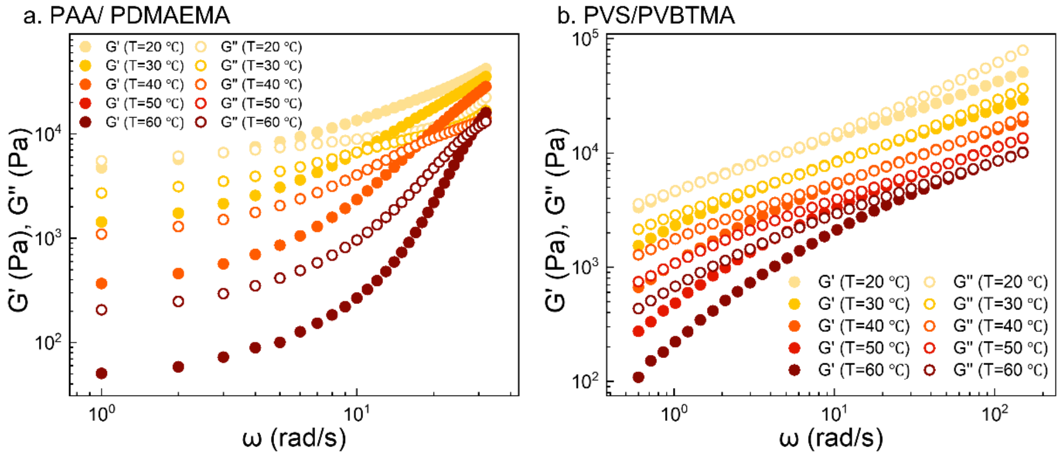


**Supplementary Fig. 10.** Frequency sweeps of (a) PAA/PDMAEMA and (b) PVS/PVBTMA at various temperatures. The storage modulus $G'$ and loss modulus $G''$ of PAA/PDMAEMA and PVS/PVBTMA coacervates monotonically decrease with the increasing temperature.

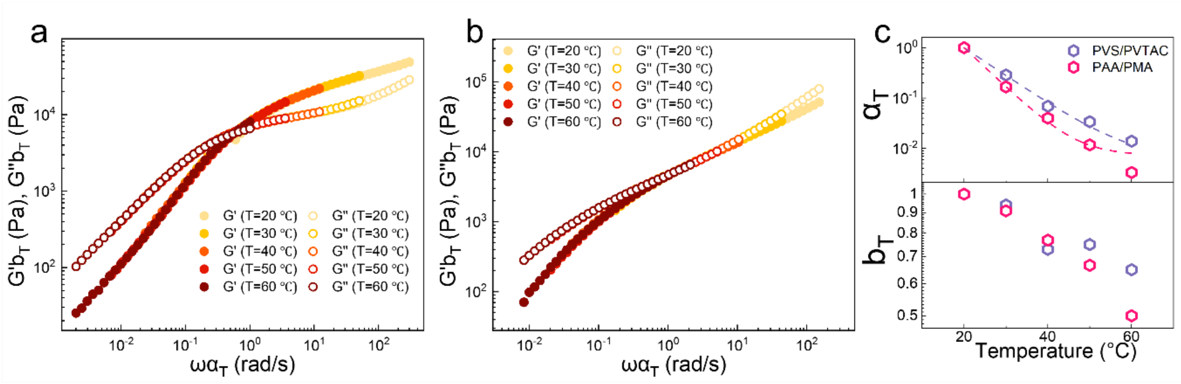


**Supplementary Fig. 11.** Rescaled frequency sweeps of (a) PAA/PDMAEMA and (b) PVS/PVBTMA at different temperatures by multiplying the frequency and modulus by horizontal and vertical shift factors, $\alpha_T$ and $b_T$, respectively. (c) Values of $\alpha_T$ and $b_T$ as a function of temperature. Experimental data of $\alpha_T$ over $T$ are fitted using the expression $\alpha = k\exp\left(E/k_B T\right)$ as indicated by the dash lines.

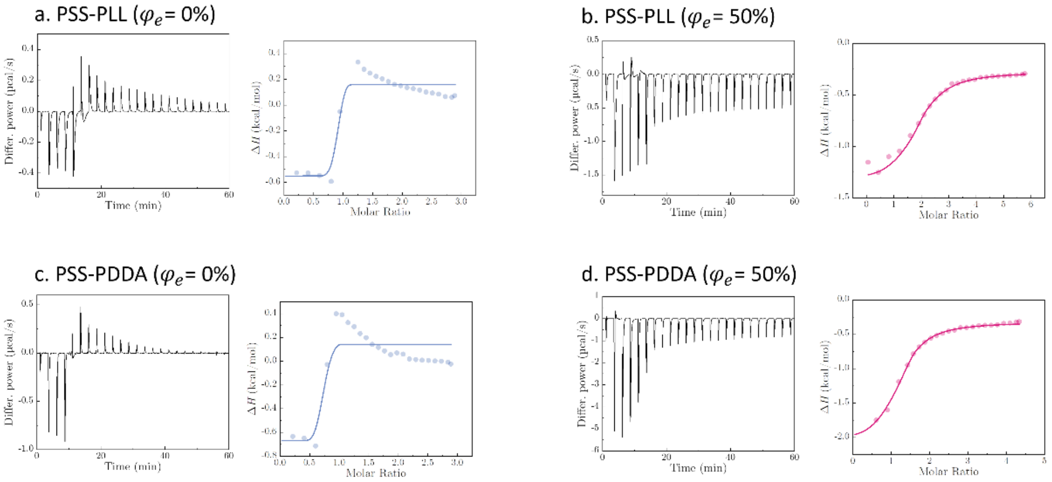


**Supplementary Fig. 12.** Plots of differential power (DP) signal versus time and the molar enthalpy change $\Delta H$ versus time PSS/PLL and PSS/PDDA coacervate at $\varphi_e = 0\%$ and $\varphi_e = 50\%$. The experimental conditions are (a) PSS (0.5 mM), PLL (7.5 mM), and $\varphi_e = 0\%$; (b) PSS (0.5 mM), PLL (15 mM), and $\varphi_e = 50\%$; (c) PSS (0.5 mM), PDDA (7.5 mM), and $\varphi_e = 0\%$; (d) PSS (0.5 mM), PDDA (11.25 mM), and $\varphi_e = 50\%$.

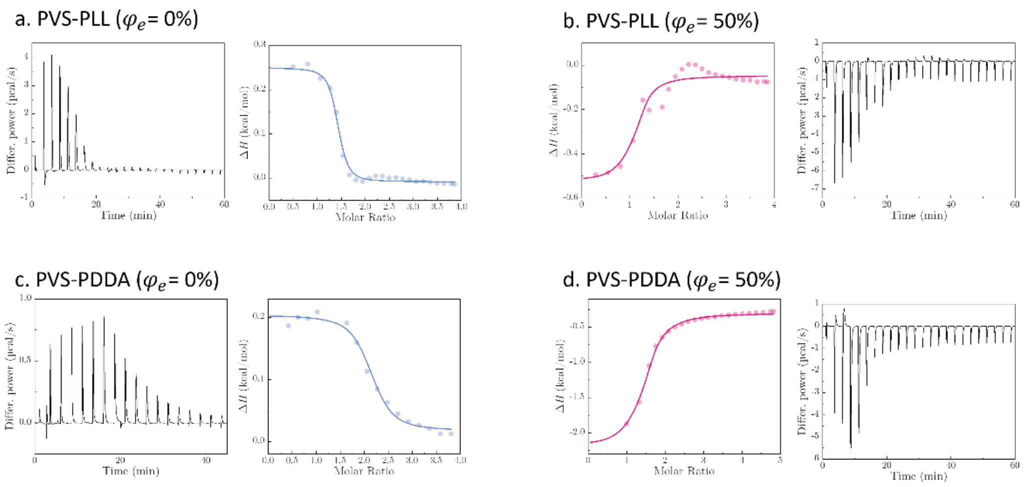


**Supplementary Fig. 13.** Plots of differential power (DP) signal versus time and the molar enthalpy change $\Delta H$ versus time PSS/PLL and PSS/PDDA coacervate at $\varphi_e = 0\%$ and $\varphi_e = 50\%$. The experimental conditions are (a) PVS (1 mM), PLL (20 mM), and $\varphi_e = 0\%$; (b) PVS (1 mM), PLL (20 mM), and $\varphi_e = 50\%$; (c) PVS (1 mM), PDDA (20 mM), and $\varphi_e = 0\%$; (d) PVS (1 mM), PDDA (25 mM), and $\varphi_e = 50\%$.

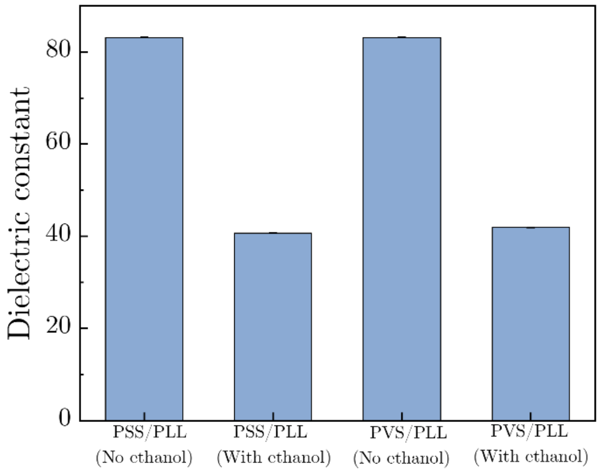


**Supplementary Fig. 14.** The computed values of dielectric constant of solutions containing oppositely charged monomers.

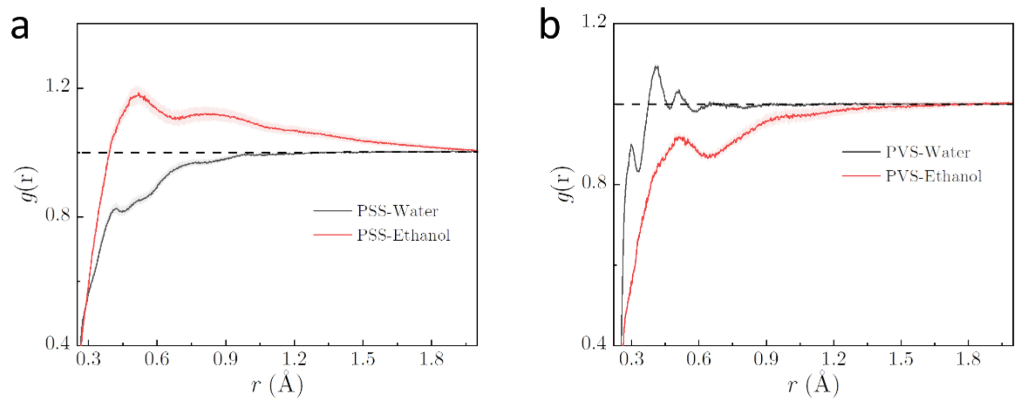


**Supplementary Fig. 15.** Radial distribution function of water (the dark lines) and ethanol (the red lines) molecules surrounding (a) PSS and (b) PVS, respectively.

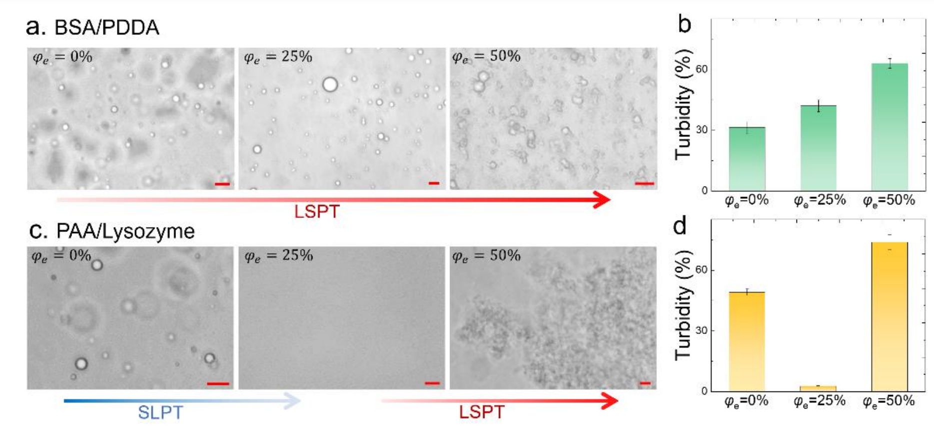


**Supplementary Fig. 16**. (a) Bright-field images and (b) solution turbidity showing the LSPT of condensates formed by BSA (2 wt%) and PDDA (1 wt%) as $\varphi_e$ increases. (c) Bright-field images and (d) solution turbidity showing SLPT and then LSPT of condensates formed by PAA (0.125 wt%) and Lysozyme (2 wt%) with increasing $\varphi_e$. Scale bars are 20 μm.

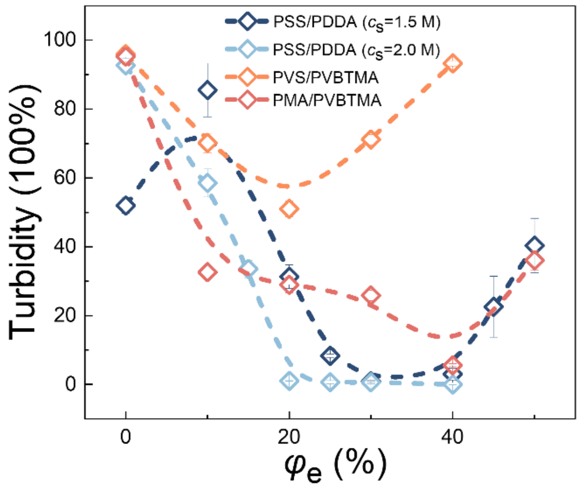


**Supplementary Fig. 17**. Solution turbidity of coacervates over the volume fraction of ethanol $\varphi_e$. These curves suggest a re-entrant phase separation of different coacervates mediated by ethanol. Coacervates are PSS (50 mM)/PDDA (50 mM) with 1.5 M NaCI, PSS (50 mM)/PDDA (50 mM) with 2.0 M NaCI, PVS(50 mM)/PVBTMA(50 mM), and PMA(50 mM)/PVBTMA(50 mM), respectively.

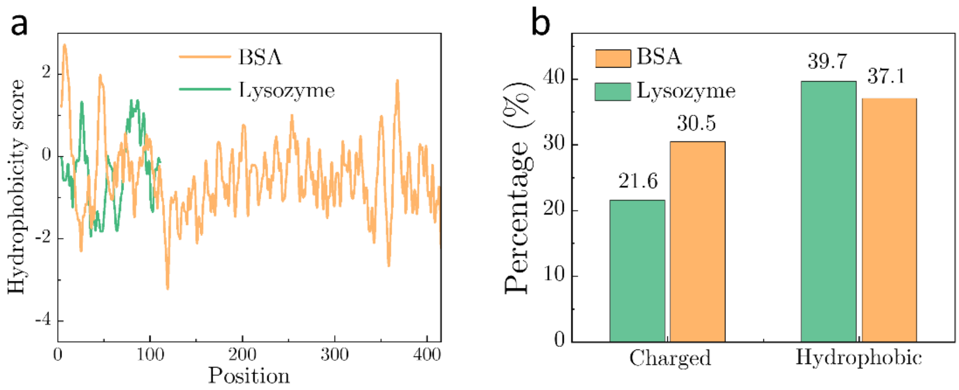


**Supplementary Fig. 18**. (a) The hydrophobicity score and (b) amino acid compositions of BSA and Lysozyme proteins.

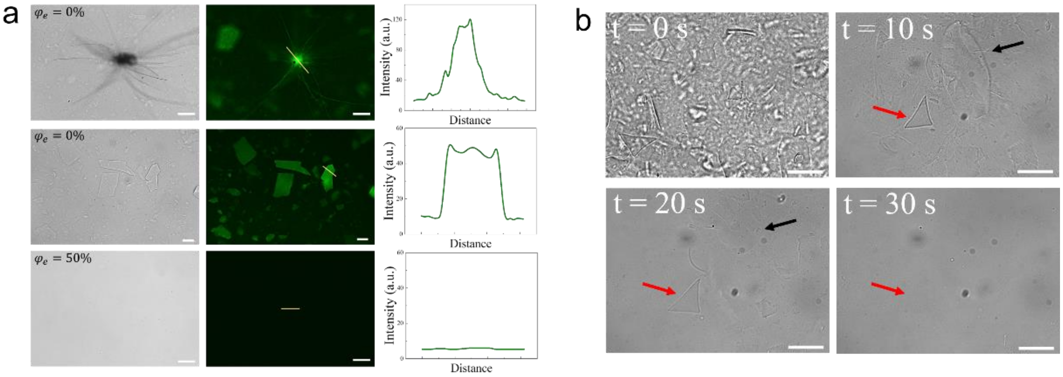


**Supplementary Fig. 19**. (a) Bright-field and fluorescence images of Aβ 1-42 plaques formed at different volume fraction of ethanol $\varphi_e$. Aβ 1-42 plaques were dyed by Thioflavin T (ThT). (b) Bright-field images showing the fast dissolution of Aβ 1-42 plaques by the introduction of ethanol at $\varphi_e = 50\%$. Scale bars are 20 μm.
